# Microbiome Analysis of the Eastern Oyster As a Function of Ploidy and Seasons

**DOI:** 10.1101/2023.08.10.552804

**Authors:** Ashish Pathak, Mario Marquez, Paul Stothard, Christian Chukwujindu, Jian-Qiang Su, Yanyan Zhou, Xin-Yuan Zhou, Charles H. Jagoe, Ashvini Chauhan

## Abstract

Shellfish, such as the eastern oysters (*Crassostrea virginica*) are not only valued as seafood but also for the ecosystem services they provide, including improving water quality and reducing eutrophication. Excess N causes eutrophication, harmful algal blooms, fish kills and overall decline of estuarine ecosystems resulting in economic losses. Oyster reefs sequester N and enhance denitrification processes, however, information on the N cycling oyster microbiome is scarce with most studies focusing on random grab samples or on pathogens, such as *Vibri*o spp. Further, triploid oysters are often used for aquaculture, as they grow faster than diploids, but there is little information on potential microbiome differences with ploidy. To address these knowledge gaps, diploid and triploid farmed oysters were collected at monthly intervals over one year and analyzed using a coupled approach encompassing shotgun metagenomics and quantitative microbial elemental cycling (QMEC) qPCR assays. Overall, the genus *Psychrobacter* dominated the core microbiome across all samples, regardless of season or ploidy, followed by *Synechococcus*, *Pseudomonas*, *Pseudoalteromonas* and *Clostridium*. *Psychrobacter* abundances increased significantly in the colder months; the same trend was also observed in the alpha and beta diversity. However, warmer months had increased bacterial diversity relative to colder months. Gene functional profiles were similar among seasons and ploidy, with respiration and metabolism of carbohydrates, RNA, and proteins as dominant functions. There were strong positive correlations between abundance of the “core” microbiome taxa and gene functions associated with central metabolism, DNA and carbohydrate metabolism, strongly suggesting the functional role of *Psychrobacter* in the microbiome. Metagenome assembly was performed to characterize dominant species, followed by phylogenetic analysis of select MAGs (metagenome-assembled genomes), further supporting the presence of multiple *Psychrobacter* spp. Sequence-based identification of denitrification genes in the *Pyschrobacter* MAGs indicated the presence of *norB*, *narH*, *narI*, *nirK*, and *norB*. QMEC analysis indicated C and N cycling genes were most abundant, with no discernable patterns due to seasons or ploidy. Among N cycling genes, the nosZII clade was dominant, which is likely responsible for the eastern oysters potential for bioextraction and enhancing water quality via denitrification.

## 1. Introduction

Oysters are consumed widely on a global scale due to their unique taste as well as nutritional value, which is high in proteins, polyunsaturated fatty acids (PUFA), vitamins and minerals (Cruz-Romero et al. 2008; Chen et al. 2019). According to estimates from 2020, U.S. oyster landings totaled 19.9 million pounds (National Marine Fisheries Service, 2020; FAO 2018; Chen et al. 2019), with an annual value of almost $187.2 million USD [(Prapaiwong et al. 2009; National Marine Fisheries Service, 2020 (https://www.st.nmfs.noaa.gov/st1/commercial/landings/annual_landings.html)]. *Crassostrea virginica*, the eastern oyster or the American oyster, accounts for as much as 75% of all commercialized oysters in the United States, with the coastal regions of Florida, Texas, Louisiana, Mississippi and Alabama, contributing to more than 60% of the total production (Prapaiwong et al. 2009).

Oysters also provide ecosystem services, including maintaining water quality and sustaining estuarine habitats (Burge et al., 2014), and thus, are recognized as keystone species across North American coasts from Florida to Maine. In fact, a single adult oyster can filter as much as 50 gallons of estuarine water per day (https://www.cbf.org/about-the-bay/more-than-just-the-bay/chesapeake-wildlife/eastern-oysters/oyster-fact-sheet.html), which is a significant bioextractive mechanism resulting in a net positive impact on water quality by way of removing particulate material and nutrients, especially the microbially-mediated removal of nitrogen (N), thus mitigating algal blooms and fish kills. In this regard, oyster aquaculture has been recognized as a long-term tool to mitigate coastal eutrophication problems (Bricker et al., 2020; Ayvazian et al., 2021), as well as an innovative approach for bioremediation of contaminants (Filippini, et al., 2022).

Despite the significance of oysters as stated above, their populations continue to decline with up to 85% of oyster habitats decimated globally, as a function of overharvesting, habitat fragmentation, decreased water quality, shifts in estuarine water salinity and diseases (Wilberg et al. 2011; Hemraj et al., 2022). According to Kirby (2004), oyster fisheries expanded and collapsed in a linear sequence along easter North America, western North America, and eastern Australia. To address this, oyster aquaculture is being used to meet the continued oyster demand, especially by applying newer genomics techniques, such as inducing polyploidy(Qin et al. 2019). Note that triploid oysters grow faster and produce higher yields relative to diploids due to partial sterility, higher heterozygosity, and different energy allocations for growth and gametogenesis (Bodenstein et al., 2023), and can be marketable year-round (Hollier, 2014). Moreover, triploid oysters appear to be less sensitive to environmental changes and stressors (Qin et al. 2019). Given these advantages, triploid oysters account for most of the commercially produced oysters in the US.

Regardless of ploidy, filter feeding behavior fosters colonization of the oyster’s nutrient-rich mucosa and digestive organs with a plethora of diverse estuarine microbiota-which are either permanently associated with the bivalve oyster, referred to as the autochthonous bacterial communities (Unzueta-Martínez et al., 2022; Romero et al., 2002; Lokmer and Wegner 2014), or can be transient, called as the allochthonous bacteria-communities that are ingested through filter feeding and simply making their way through the bivalves gills and digestive organs. The autochthonous “core” microbiota can forge a mutually beneficial symbiotic relationship with their bivalve host organism; however, a universal “core” pertaining to host-microbe interactions remains unclear (Shade and Handelsman, 2012). Furthermore, it also remains to be known whether the triploid oysters, due to their larger size, filter a higher biomass of bacteria from the water column and accumulate larger concentrations and/ or diversity of beneficial or pathogenic mortality inducing bacteria, thus represented with a different ‘core” microbiome compared to the diploids. To this end, De Deckar et al. (2011) reported a higher susceptibility of triploid oysters to the pathogenic *Vibrio* bacteria relative to their diploid counterparts.

It was Colwell and Liston that first suggested the existence of a defined commensal flora in oysters dating back to 1960. Zurel et al. (2011) have shown a conserved seasonal association between the Chama-associated oceanospirillales group (CAOG) of bacteria with oysters, likely representing a symbiotic association. Trabal et al. (2012) reported symbiotic host-bacteria relationships during different growth phases of two oyster species- *Crassostrea gigas* and *Crassostrea corteziensis*. Such symbionts may assist in the digestion processes, as has been demonstrated in the larvae of *Crassostrea gigas* (Prieur et al., 1990), and may also supply the bivalve host with vitamins and amino acids that serve as growth factors-as shown in the Pacific vesicomyid clam-*Calyptogena magnifica* (Newton et al., 2007). Moreover, certain symbiotic bacteria can even protect their host from pathogens by either producing antimicrobial agents, or by growing in high densities that prevents colonization by other strains (Pujalte et al., 1999). More recently, Sakowski et al., 2020 discovered that up to 33% of the microbiota from the extrapallial fluid, responsible for shell formation, were autochthonous to *C. virginica*. Interestingly, the extrapallial fluid-associated microbiota were reported to be rich in functions related to dissimilatory nitrate reduction, nitrogen fixation, nitrification, and sulfite reduction. Banker and Vermeij (2018) reported that sulfate reduction bacteria (SRB) play significant roles in calcification, but they may not be the only bacteria involved in this process. Moreover, the sulfate and nitrate reduction were proposed to have a synergistic effect on calcium carbonate precipitation and thus likely impacting shell formation (Sakowski et al., 2020). It was also previously shown by Braissant et al. (2007) that SRBs can produce exopolymeric substances (EPS) which can serve as a site for CaCO_3_ precipitation, and that the calcifying ability can be enhanced by an increase in alkalinity due to the reduction in sulfate ions. Furthermore, most calcifying bacterial are also actively involved in nitrate reduction (Cacchio et al., 2012). In this regard, beneficial oyster microbiota likely enhance their host health, thus promoting growth and longevity. To further tease out the transient (allochthonous) oyster microbiota from the resident ones, a flexibility score (FS) was recently developed, which showed that the gill tissues harbor a diverse range of resident bacteria (Unzueta-Martínez et al., 2022).

The major bacterial phyla of the eastern oysters are as follows: Proteobacteria, Cyanobacteria, Bacteroidia, Mollicutes, Bacteroidetes, Tenericutes and Firmicutes (Hines et al., 2023; Akter et al., 2023; Singh et al., 2023; Sakowski et al., 2020; Pierce et al., 2019; King et al., 2012; Arfken et al., 2017; Stevick et al., 2019; Pimentel et al., 2021; Banker and Vermeij 2018). A consensus on the genus level “core” and possibly the resident or autochthonous microbiome of eastern oysters is lacking, but the following bacteria, in addition to *Vibrio* species, have been identified to be dominant groups: *Mycoplasma*, *Pseudoalteromonas*, *Burkholderia*, *Bacteroides*, *Lactobacillis*, *Acetobacter*, *Allobaculum*, *Ruminococcus*, *Nocardia,* and *Oceanospirillales* (Britt et al., 2020; Unzueta-Martínez et al., 2022). It is noteworthy that the bacterial communities within the eastern oyster and its specific niches, such as the mantle fluid, gills, digestive system, gut, extrapallial fluid, and hemolymph have been found to be distinctly different than the surrounding water or sediment communities (Akter et al., 2023; Unzueta-Martínez et al., 2022; Sakowski et al., 2020; Lokmer and Wegner 2015; King et al. 2012; Wegner et al. 2013). In one previous study conducted on the wild type and farmed eastern oysters, we found Cyanobacteria accounting for 50-75% of the total microbial communities, based on 16S rRNA amplicon metagenomics (Chauhan, et al., 2014). In another microcosm-based study related to the Deepwater Horizon oil spill, *Pseudomonas* spp. were found to be the dominant oyster-associated bacteria (Thomas et al., 2014; Chauhan et al., 2014; Pathak et al., 2021; Chauhan, et al., 2013). More recently, our metagenomics survey of the eEastern oysters indicated dominance of *Lactobacillus, Burkholderia*, *Bradyrhizobium*, *Afipia* and *Delfitia* bacterial groups with lower representations of *Psychrobacter* spp. Additionally, *Mycoplasma*, which has been previously suggested as one of the autochthonous bacteria in eastern oysters, was also identified (unpublished data).

One major ecosystem service associated with the oysters relates to enhancing water quality by utilization of the dissolved nitrogen by both oysters and their microbial communities. Specifically, N assimilated into oyster shells and tissues is removed from the estuarine systems via harvesting or burial (Vitousek et al., 1997). Oyster feces and pseudofeces contribute organic material to sediments, potentially enhancing dentrification processes. Denitrification is a four-step reduction of nitrate (NO^3-^) to N_2_ via nitrite (NO2^-^), NO and N_2_O and eventually to nitrogen gas (Ray et al., 2021; Ayvazian et al., 2021). At the genetic level, denitrifying bacteria harbor genes in operonic clusters that encode functions for reduction of nitrate (narG/napA), nitrite (nirS/nirK), nitric oxide (norB/norC), and nitrous oxide (nosZ) (Zumft, 1997).

This led us to the following critical question - what are those suits of environmentally relevant biogeochemical functions, such as nitrogen cycling, that are performed by the eastern-oyster associated microbiome communities? To address this, we applied shallow shotgun sequencing (SSS), recognized as a viable and cost-effective method to conduct both taxonomic and functional gene analysis in complex environmental samples (Lugli et al., 2022). To augment the findings obtained from SSS we applied the recently developed method of “Quantitative Microbial Elemental Cycling (QMEC)”, a high-throughput quantitative PCR (HT-qPCR) technique for estimating microbial gene copy numbers engaged in critical steps of Carbon (C), Nitrogen (N), Phosphorus (P), and Sulfur (S) cycling as well as methane metabolism (Zheng et al., 2018). Given that the commercially grown eastern oysters are triploid variants that originated from the wild-type diploids, and that their microbially-mediated functions likely oscillate on a continuum, both diploid and triploid eastern oysters were collected monthly over an annual growth cycle to: 1) conduct shotgun metagenomics and delineate native autochthonous microbial groups; 2) evaluate the microbial gene functional shifts occurring within the microbiome ; and 3) evaluate microbially-mediated CNPS biogeochemical cycling by applying the QMEC method.

## 2. Materials and Methods

### 2.1. Sample Collection and Environmental Measurements

Cultured *C. virginica* were collected from the aquaculture research lease of the Wakulla Environmental Institute located in Oyster Bay [30.0342702, -84.3558844], near Panacea, Florida. Oysters were grown in plastic cages, suspended from lines attached to pilings at depths of 1-2 meters (Australian Adjustable Long-Line System; https://seapa.com.au/farming-systems/). At each sampling period, 15 diploid and 15 triploid oysters were measured using digital calipers. Shell height was measured as the maximum distance from umbo (hinge) to lip (commissure) (per Galtsoff, P. S. 1964. The American Oyster *Crassostrea virginica* Gmelin. Fishery Bulletin, v. 64. United States Government Printing Office, Washington, D. C.) Over an annual growth period spanning from December 2016-August 2017, diploid and triploid oysters were collected, stored in Ziploc bags over ice, returned to the laboratory and stored at -80 °C until further analysis.

Temperature and salinity were measured during weekly visits to the oyster farm using a YSI Model 85 (YSI Instruments, Yellow Springs OH) water quality meter equipped with a salinity/conductivity/temperature/dissolved oxygen sensor probe calibrated according to the manufacturer’s directions. Monthly water samples were collected from approximately 20 cm below the surface into acid-cleaned HDPE bottles (Nalgene), placed on ice, and returned to the laboratory for filtering.

Total suspended solids and suspended organic matter were determined after filtration by drying and loss on ignition, respectively (Standard Methods for the Examination of Water and Wastewater, https://www.wef.org/publications/publications/books/StandardMethods.) One liter samples were filtered through tared Whatman GF/F (borosilicate glass fiber filters, pore size 0.7 µM) using a glass vacuum filtration flask and funnel. Filters were dried to constant weight at 60° C in a Fisher Isotemp drying oven, then ashed at 55° C in a Thermoline 48000 muffle furnace and reweighed. Filtered water samples were aliquoted into acid washed HDPE bottles and frozen at -20° C. Chlorophyll a was measured per the EPA Standard Method LG405 (Revision 09, March 2013), following the original method (Welschmeyer, 1994).

### 2.2. DNA Extraction and Library Preparation for Next-Generation Sequencing (NGS)

Each oyster was first externally rinsed with sterile water and scrubbed thoroughly to remove the shell biofilm, bivalves were then opened with a shucking knife at the hinge where the adductor muscle is located joining the two shells. Once opened, the oyster’s mantle fluid was drained, and the entire oyster was collected. Pooled (n=10) diploid and triploid oysters from each time point were then homogenized thoroughly using the Oster brand kitchen blender and processed for DNA extraction.

At each time, 250 mg of homogenized oyster tissues were used to extract DNA in duplicate, labeled as A and B, using the Qiagen MagAttract PowerSoil DNA KF kit (Formerly MOBio PowerSoil DNA Kit). DNA quality was evaluated visually via gel electrophoresis and quantified using a Qubit 3.0 fluorometer (Thermo-Fischer, Waltham, MA, USA). Libraries were then prepared by MicrobiomeInsights Inc. (https://microbiomeinsights.com/) using an Illumina Nextera library preparation kit using their in-house protocol (Illumina, San Diego, CA, USA).

### 2.3. DNA Sequencing, Quality Control Analysis, Data Curation, and Sequence Processing

Paired-end sequencing (150 bp x 2) was done on a NextSeq 500 in medium-output mode. Shotgun metagenomic sequence reads were processed with the Sunbeam pipeline. Initial quality evaluation was done using FastQC v0.11.5 ( https://www.bioinformatics.babraham.ac.uk/projects/fastqc/) followed by adapter removal, read trimming, low-complexity-reads removal, and host-sequence removals. Adapter removal was done using cutadapt v2.6 (Martin 2011) andtrimming was done with Trimmomatic v0.36 (Bolger et. al., 2014) using custom parameters (leading 3, trailing 3, sliding window 4:15, minimum length 36). Low-quality sequences were detected with Komplexity v0.3.6 (Clarke et al. 2019) and discarded from further analysis. High-quality reads were mapped to the human genome (Genome Reference Consortium Human Reference 37) and to the *Crassostrea virginica* genome v3.0 (Accession number GCF_002022765.2); matching were removed (labeled as host in Fig. SI-1), along with low quality reads (marked as low quality in Fig. SI-1), respectively. Among the high-quality reads, six sequenced samples were considered outliers because they contained variable number of host reads. These samples, represented by n=10 pooled oysters included triploid oysters from January (010717 Triploid A and 010717 Triploid B), Diploid B oysters from May (051117), Diploid A oysters from December 2016 (120516), and both duplicate triploids from December 2016 (120516 Triploid A, and 120516 Triploid B), respectively (note that samples are labeled based on their collection date and ploidy). To retain the high quality read samples, we subsampled five of them to the median value of reads for the non-outliers (6768 reads). Sample 051117 Diploid B had a similar number of reads to the other samples, so it was not subsampled. The remaining reads, marked as retained in Fig. SI-1, were taxonomically classified using Kraken2 (Wood et al. 2019) with the minikraken v1 database. Functional profiles were not subsampled, and the five outliers were removed from the downstream bioinformatics analysis. For functional profiling, high-quality (filtered) reads were aligned against the SEED database via translated homology search and annotated to subsystems, or functional levels, 1-3 using Super-Focus (Silva et al. 2016).

### 2.4. HT-qPCR Quantitative Microbial Element Cycling (QMEC)

Based on high-throughput qPCR (HT-qPCR) , a recently developed method called QMEC [Quantitative Microbial Elemental Cycling (QMEC)], has the potential to simultaneously detect and quantify biogeochemical cycling genes for C, N, P and S, thus. providing a deeper understanding on these microbially-mediated processes (Zheng et al., 2018). Briefly, a total of 72 primer sets were used to quantify DNA extracted from homogenized oyster samples. The primer sets targeted the 16S ribosomal RNA (rRNA) gene, and 35 carbon cycling genes (C), 22 nitrogen cycling genes (N), 9 phosphorus cycling genes (P) and 5 sulfur cycling genes (S). Amplification for each sample was conducted in triplicates, using 100 nL containing (final concentration) 1 × Light Cycler 480 SYBR® Green I Master Mix (Roche Inc., USA), nuclease free PCR-grade water, 1-3 ng μL DNA template, and 1 μM each of forward and reverse primers. High-throughput qPCR (HT-qPCR) was performed by the Smart Chip Real-time PCR system (Takara Bio USA, Inc.). A non-template negative control was set for each primer set. The thermal cycle curve conditions included an initial denaturation at 95 °C for 10 min with 40 cycles of denaturation at 95°C for 30 s, annealing at 58°C for 30 s and extension at 72°C for 30s. The melting curve analysis was automatically generated by the software, and the qPCR results were analyzed using Smart Chip qPCR software. Samples with efficiencies beyond the range 1.7–2.3 or an r^2^ under 0.99 were discarded. A threshold cycle (Ct) 31 was used as the detection limit. All three replicates that amplified successfully were used for further data analysis. The relative gene copy number was calculated as described by Looft et al., 2012 using following equation.

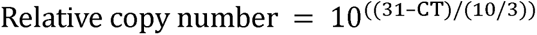

where CT refers to HT-qPCR results, 31 refers to the detection limit.

Absolute 16S rRNA copy numbers were determined by the standard curve (SC) method of quantification using the Roche 480 system. Each 20 µl qPCR consisted of 10 µl 2× Light Cycle 480 SYBR Green I Master (Roche Applied Sciences), 1 µM each primer, 2-6 ng DNA as template and enough nuclease-free PCR-grade water. The thermal cycle consisted of 5 min initial enzyme activation at 95°C, followed by 40 cycles of denaturation at 95 °C for 30 s, annealing at 60 °C for 30 s and extension 72 °C for 20 s. A plasmid control containing a cloned and sequenced 16S rRNA gene fragment (6.69 × 10^8^ copies per microliter) was used to generate five-point calibration curves from tenfold dilutions. All qPCRs were performed in technical triplicates with non-template reactions serving as negative control.

### 2.5. Summary of Data Transformation

All qPCR data was normalized into relative and absolute abundance. The relative abundance was calculated by normalizing the relative gene copy of each C N, P, and S-cycling genes with its relative 16S copy number (equation I) (Zheng et al., 2018). The absolute abundance was calculated by multiplying the 16S copy number with the relative abundance of each C, N, P, and S-cycling genes and dividing the product with its respective DNA concentration (equation II) (Zheng et al., 2018). The calculated absolute abundance was then normalized into copies/g oyster among samples (equation III) (Zheng et al., 2018), which was used to evaluate the total CNPS genes in each sample.

### 2.6. Statistical Analysis

Averages and standard deviations were calculated in Microsoft Excel. Statistical visualizations were done in R and Origin statistical software. Heatmaps, correlation, hierarchical clustering analysis and were done on R studio (https://www.R-project.org/) using pheatmap, vegan, corrplot and ggplot 2. The stacked bar charts were plotted in Origin (https://www.originlab.com/).

Negative binomial models (DESEq2 R package) were used for differential abundance testing of taxonomic and subsystem level features. Changes due to sampling timepoint differences after controlling for genotype differences (ploidy) were analyzed; because we did not notice major differences between genotypes by PERMANOVA, we used all samples together even though several timepoints were missing for the diploid samples. We plotted the top 20 groups for each test, based on their adjusted p-values, which were calculated using the likelihood ratio test (LRT). The critical p-value (alpha value) were used to compare the adjusted p-values (= 0.01) of the multiple independent groups, unless otherwise stated.

Furthermore, abundance of the top 10 bacterial taxa and gene functions were averaged to produce a single dataset for each sampling time point representing diploids or triploid samples which was then imported into the Primer-E software suite (version 6.1.18; PRIMER-E, Ivybridge, United Kingdom). Normalization of data was performed using the log (X+1) pre-treatment function in the Primer-E software and after transformation, a Bray-Curtis similarity matrix was generated, and non-metric multidimensional scaling plot (NMDS) was obtained using parameters of Kruskal stress formula1 and minimum stress 0.01. Dendrogram analysis was performed based on the group average option using Primer-E software.

### 2.7. Shallow Shotgun Metagenome Submission: Metagenomic Sequence Accession Numbers

The shallow shotgun metagenomic sequences obtained from this study are available under NCBI accession # PRJNA858118, Biosample #SAMN29671270 to SAMN29671291 and SRA#SRS13888819 to SRS13888833(https://www.ncbi.nlm.nih.gov/bioproject/PRJNA858118/).

### 2.8. Metagenome Assembly

Metagenomes were assembled, binned into MAGs (metagenome-assembled genomes), and annotated using the nf-core/mag pipeline v2.1.1 (Krakau et al., 2022). Raw reads from 22 samples were used as input. Reads were trimmed using fastp v0.20.1 (Chen et al., 2018) and assembled using SPAdes v3.15.3 (Prjibelski et al., 2020). Metagenomic contigs were grouped into draft genomes (i.e., binned into MAGs) using MetaBAT 2 v2.15 (Kang et al., 2019). Bin quality was assessed using BUSCO v5.1.0 (Simão et al., 2015) and Quast v5.0.2 (Gurevich et al., 2013). Bins with a level of completeness less than 15% (based on BUSCO results) were excluded from further analysis. Taxonomic classification of bins was performed using GTDB-Tk v1.5.0 (Chaumeil et al., 2019). Bins were annotated using Prokka v1.14.6 (Seemann, 2014).

### 2.9. Phylogenetic Analysis of Binned Metagenomes

Prokka-derived protein sequences from four metagenomic bins classified as *Psychrobacter* were used to construct a phylogenetic tree, using PhyloPhlAn v3.0.67 (Asnicar et al., 2020). 215 RefSeq *Psychrobacter* proteomes available in the NCBI Assembly database were downloaded and included in the analysis, as well as a *Moraxella lincolnii* protein set (GCF_002014765.1), for use as an outgroup. The “phylophlan” database of 400 universal marker genes was used, and a diversity setting of “medium” was specified. RAxML v 8.2.12 (Stamatakis, 2014) was used to generate a maximum likelihood tree with bootstrap values from 100 replicates.

### 2.10. Average Nucleotide Identity (ANI) Analysis of MAGs

Whole-genome similarity between each *Psychrobacter* MAG and all NCBI RefSeq *Psychrobacter* genomes was assessed by ANI, calculated using FastANI v1.33 (Jain et al., 2018). For the ANI analysis all RefSeq *Pyschrobacter* assemblies were compared to each MAG. A separate ANI result was generated for each comparison, providing a total of 993 ANI values. For each MAG the highest ANI was identified and that RefSeq sequence is reported. No minimum ANI threshold was used in this process.

### 2.11. Identification of Denitrification Genes in MAGs and RefSeq Genomes

Prokka-derived proteins for *Psychrobacter* MAGs and protein sequences from NCBI for similar *Psychrobacter* RefSeq genomes (based on ANI) were analyzed for denitrification genes using NCycDB (Tu et al., 2019). Searching was performed against the NCyc_100.faa database using DIAMOND v2.0.14.152 (Buchfink et al., 2021). The following genes / proteins were assessed: *napA* (Periplasmic nitrate reductase NapA), *napB* (Cytochrome c-type protein NapB), *narG* (Nitrate reductase), *narH* (Nitrate reductase), *narI* (Nitrate reductase gamma subunit), *nirK* (Nitrite reductase), *nirS* (Nitrite reductase), *norB* (Nitric oxide reductase subunit B), *norC* (Nitric oxide reductase subunit C), *nosZ* (Nitrous-oxide reductase). NCycDB results were compared to Prokka and NCBI annotations, and additional BLAST searches against the NCBI RefSeq and UniProtKB/Swiss-Prot databases were performed to resolve conflicting functional assignments.

## 3 Results and Discussion

### 3.1. Site Characterization and Shallow Shotgun Sequencing (SSS)-based Metagenomics of the Eastern-Oyster Over an Annual Growth Cycle

At the start of the growing period, diploid oysters ranged from 21-35 mm in height and 1-3 g in weight; triploid oysters heights were 12-27 mm and weights were 0.5-1 g. After one year of growth, diploids were 85-95 mm in height and 54-91 g in weight, and triploids were 71-105 mm in height and 75-110 g in weight (cultured oysters are considered harvestable when height > 76 mm). Water depths in the area ranged from 0.5-3.5 meters, depending on tides, and the water column was well mixed by winds and currents, with little or no stratification with depth. Over the annual sampling time-period (December 2016-August 2017), water temperatures ranged from 13-31° C, with a median of 26.5 °C. Salinity ranged from 16.1-28.3 psu, with a median of 23.5 psu. Total dissolved nitrogen ranged from 35.7-1000.1 μM (median 57.1 μM). Particulate organic matter ranged from 3.3-29.6 μg/L (median 10.8 μg/L) and chlorophyll-a ranged from 0.69-25.33 μg/L (median 6.93 μg/L).

For the metagenomic analysis, 10 diploid and 10 triploid oysters were collected at each monthly time point, thus a total of 240 oyster samples were collected. Shallow shotgun sequences (SSS) obtained from n=10 each of diploid and triploid oysters, collected in duplicates (labeled as A or B), yielded over 113.2 million raw reads and 98.3 quality-filtered reads (Fig. SI-1). Around 34.3 (34.8%) and 0.5 million (0.51%) of the quality-filtered reads were assigned to the oyster host (*Crassostrea virginica* genome v3.0 (Accession number GCF_002022765.2) and the human genome (Genome Reference Consortium Human Reference 37), which were removed prior to further processing.

One goal of this study was to examine the eastern oyster microbiome over an annual growth cycle with oysters collected every month, rather than using random grab samples, which are appropriate only to provide a quick snapshot of bacterial communities but are not sensitive in the assessment of the eastern oyster’s autochthonous microbiome communities. Once the “core” microbiome of the Eastern oyster is better understood, specific studies can be conducted to identify those microbial groups that are specific to beneficial functions such as maintenance of oyster growth, overall health, and productivity. To this end,regardless of seasons or ploidy, the eastern oysters were found to be predominantly colonized by bacteria, as shown by 93.4% of sequences that were annotated to the bacterial domain, followed by eukaryotes (5.64%), archaea (0.76%) and viruses (0.25%), respectively. Among bacteria, a total of 36 phyla were identified across all the samples with the top 5 phyla represented by *Pseudomonadota* (58%; former *Proteobacteria)*; *Bacillota (11%; former Firmicutes*; *Bacteroidota (7%; former Bacteroidetes)*; *Cyanobacteria* (6%) and *Actinomycetota (3%; former* Actinobacteria), respectively. *Pseudomonadota* are metabolically and ecologically diverse and are commonly found in many environments, including animal hosts; some proteobacterial groups are also pathogenic. These findings are in line with other previous studies, which show the dominance of proteobacterial lineages in oysters (Chauhan et al. 2014; Kobiyama et al. 2018; Yu et al. 2021; Pimentel et al. 2021; Sakowski et al., 2020; Pierce and Ward, 2019; Li et al., 2022). *Pseudomonadota* also likely play important functional roles in the eastern oysters, such as nitrogen cycling (Arfken et al., 2017; Ray et al., 2021; Ayvazian et al., 2021). Furthermore, proteobacterial members can also metabolize cellulose and agar-major components of the oyster’s phytoplanktonic diet, and hence colonize bivalve gastrointestinal tracts (Trabal et al., 2014), providing nutrients and bioactive growth factors to their oyster hosts (Kamiyama et al. 2009; Natrah et al. 2014; Stevick et al. 2019; Romero et al., 2002; Hernández-Zárate and Olmos-Soto, 2006; Zurel et al., 2011).

Among proteobacterial members, the gamma-proteobacterial class was the most abundant, as also shown by (Pierce and Ward 2019), followed by alpha-and delta proteobacteria. These proteobacterial groups are typically found in high numbers in marine environments, and are concentrated in oysters by filter-feeding estuarine water (Rappé et al. 2000; Kersters et al. 2006; Trabal et al. 2014; Pierce and Ward 2019). Also, gamma-proteobacteria have been frequently detected in both healthy or diseased shellfish including oysters (Pujalte et al. 1999) and as endosymbionts from the gills or gonads of different bivalve species (Dubé et al., 2019; Melissa et al., 2018). Therefore, gamma-proteobacteria are likely to be autochthonous to oysters (Paul et al., 2022).

As shown in Fig. 1A, the top 3 gamma-proteobacterial genera included *Psychrobacter* (34%); *Pseudomonas* (3.2%); and *Pseudoalteromonas* (2.7%). Besides these gamma-proteobacteria, cyanobacterial *Synechococcus* spp., from class Cyanophyceae was the second most abundant bacteria (3.6%). These bacterial genera likely constitute the “core” microbiomes in the eastern oysters analyzed in this study, using the MicrobiomeAnalyst pipeline, at threshold of 10% sample prevalence and 0.01% relative abundance (Fig. 1B). The top 10 bacterial “core” genera were *Psychrobacter*, *Synechococcus*, *Pseudomonas*, *Clostridium*, *Pseudoalteromonas*, *Mycoplasma*, *Bacillus*, *Staphylococcus*, *Vibrio* and *Cyanobium*, (Fig. 1B). *Psychrobacter* represented 6 out of the dominant 10 species found in these eastern oysters, including *P.* sp. P11F6; *P*. sp. 28M-43; *P*. sp. P11G3 (data not shown).

**Fig. 1.**
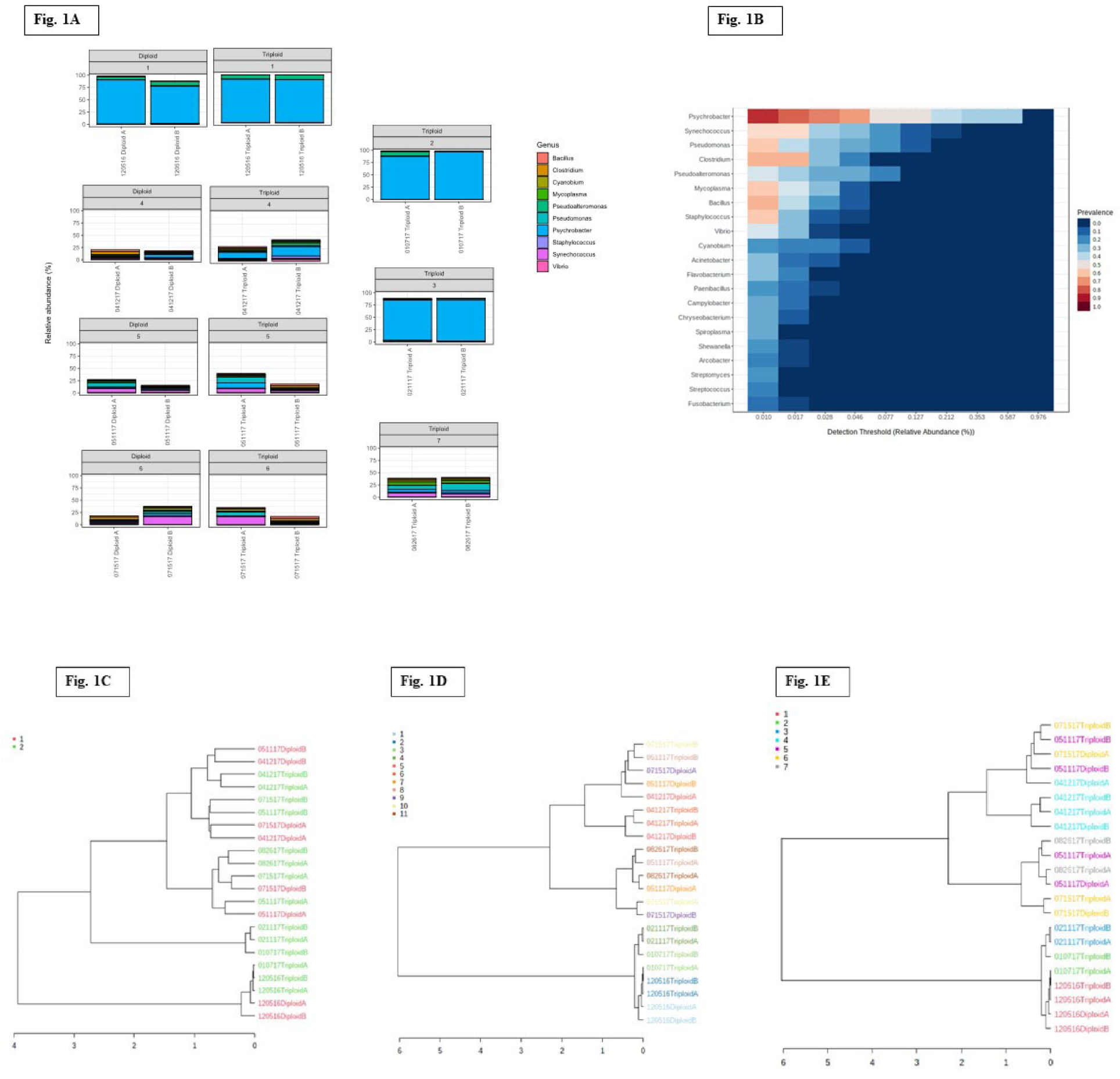
Shown are the bacterial genera-level taxonomic groups identified in diploid eastern oyster vs. triploids, sampled in duplicates (each time point is labeled as A or B), over an annual cycle. Shown are A, metagenome sequences grouped as originating from either diploid or triploid oysters, regardless of sampled time; B, core microbiomes identified in the eastern oyster, regardless of ploidy or sampled seasons; C, dendrogram analysis conducted on bacterial genus-level data obtained from diploid vs. triploid populations with group 1 containing diploids and group 2 containing triploids; D, dendrogram analysis conducted on samples binned as a function of sampled time period and ploidy was ignored; E, dendrogram analysis conducted on sample grouped as a function of collection time period: group 1 (December 2016); group 2 (January, 2017); group 3 (February, 2017); group 4 (April, 2017); group 5 (May, 2017); group 6 (July, 2017); group 7 (August, 2017), respectively.

*Psychrobacter* are known to typically thrive in colder marine ecosystems (Bowman, 2006), such as glacial ice (Zeng et al., 2016), sea ice (Bowman et al., 1997), permafrost (Bakermans et al., 2006), as well as found associated with deep sea animal host species e.g., sponge, porpoise, crab, krill, seal, and cyanobacterial mats (Welter et al., 2021; Apprill et al., 2017; Kämpfer et al., 2020; Bakermans, 2018). However, warmer niches have also been found to be colonized by *Psychrobacter* spp. including humans, lambs, and fecal materials from pigeons (Kampfer et al., 2002; Welter et al., 2021). In a recent genomic and phenotypic comparison of an extensive cohort of *Psychrobacter* spp. Welter et al., (2021) showed that the *Psychrobacter* can be delineated into two growth profiles: the “flexible ecotype” (FE) group that could grow in temperatures between 4-37°C, and the “restricted ecotype” (RE) group that grew between 4-25°C temperature. The dominant *Psychrobacter* spp. in this study are closely related to those isolated from marine Arctic environments, indicating that the eastern oyster surveyed in this study likely harbor the RE *Psychrobacter* ecotypes. Proliferation of psychrotrophic bacteria that create conditions for seafood spoilage by hydrolysis of proteins and lipids in oyster tissues, producing malodorous compounds and toxic byproducts (Madigan et al. 2014), can be a cause for concern. Beneficial probiotic properties of *Psychrobacter* have also been reported in some studies, such as enhancing growth and survival of shrimp larvae (Franco et al. 2016), and by enhancing autochthonous microbiomes and protection from bacterial pathogens (e.g., *Vibrio* sp.) in Atlantic cod (Caipang et al. 2020; Makled et al., 2017; Ramírez et al., 2020). Therefore, it may be possible that *Psychrobacter* spp. are beneficial to eastern oysters in some similar manner. Interestingly, it has been reported that oyster processing, including refrigeration, storage, and depuration at 10°C, caused a shift in the predominant *Vibrio* species that are characteristic of freshly harvested oysters, towards psychrotrophic bacteria consisting of *Pseudomonas* spp., *Aeromonas* spp., *Shewanella* spp., and *Psychrobacter* spp. (Tokarskyy et al., 2019). Therefore, further research on the dominance of *Psychrobacter* species in eastern oysters should be conducted to evaluate whether these species provide benefit(s) to their oyster host or represent an underexplored agent of oyster spoilage and pathogenicity.

In this regard, some species of *Psychrobacter,* mainly *P*. *sanguinis*, *P*. *phenylpyruvicus*, *P*. *faecalis*, and *P*. *pulmonis* are known to cause opportunistic infections in mammals (Deschaght et al., 2012; Wirth et al., 2102), which needs to be further investigated. Human infections from consuming raw or undercooked shellfish containing a higher abundance of *Psychrobacter* species has not been demonstrated, to our knowledge but neither has the predominance of *Psychrobacter* been shown for eastern oysters prior to this work. Given that the *Psychrobacter* identified in eastern oysters using metagenomics in our study appear mainly to be the RE ecotypes, it is likely that they can mount mammalian infections that require warmer temperatures, but this aspect remains unclear. As stated before, genomic analysis has recently suggested that *Psychrobacter* ecotypes likely evolved from a mesophilic common ancestor, which presents a unique opportunity to delineate the evolutionary mechanisms of the eastern oyster associated *Psychrobacter* species as a function of their estuarine habitats.

When compared over an annual cycle, a decline in bacterial abundances in eastern oysters was observed in samples that were collected from April 12, 2017, to August 26, 2017. For example, proteobacterial abundances were predominant at an average of 97% between December 2016 to February 2017), but declined to 36% from April to August 2017. Along these lines, it was recently shown for Australian oysters and mussels that temperature strongly correlated to their microbiomes and that microbial diversity increased in winter relative to summer timepoint (Akter, et al., 2023). However, low temperature likely favored the observed increase of the psychrotrophic *Psychrobacter* species (Fig. 1B). Interestingly, dendrogram analysis conducted on bacterial metagenomes revealed less affiliation with regards to ploidy; conversely, seasonal changes appeared to have a stronger influence in structuring the oyster microbiome (Fig. 1C-E). Specifically, when metagenomes were grouped as originating from either diploid or triploid oysters (Fig. 1C), a total of 6 clusters representing different time periods were observed. The general trend was that the metagenomes from the colder months-December, January and February, clustered together, as did those from the warmer months of April, May, July and August. When sequences were grouped as a function of sampled time and ploidy was ignored, 5 clusters emerged, and again, the season exerted a stronger influence in driving the bacterial communities, relative to ploidy (Fig. 1D). As a third option, when samples were grouped as a function of the sampled time, 6 clusters were observed which clearly shows seasonal impacts to be in the major factor structuring the oyster-associated microbiomes, when compared with ploidy (Fig. 1E). This finding is noteworthy because most studies on the eastern oysters have been conducted using samples collected at random times during the year (Singh et al., 2023; Sakowski et al., 2020; Pierce et al., 2019; King et al., 2012; Arfken et al., 2017; Stevick et al., 2019; Pimentel et al., 2021), and even those that were samples on a spatio-temporal basis focused on pathogen dynamics such as *Vibrio* and their occurrence in the eastern oysters.

The other two main bacterial genera identified were *Synechococcus* and *Pseudomonas* spp. (Fig. 1A and B). *Synechococcus* spp. are commonly found in estuarine waters and have also been reported from the digestive gland *C*. *gigas*, as well as connective tissue, mantle, and gonad of oysters (Avila-Poveda et al., 2014). Our previous studies along with several others have demonstrated pseudomonads to be a major bacterial group in the oysters (Chauhan et al., 2013; Thomas et al., 2014; Pathak et al., 2021). Notably, some pseudomonads can be opportunistic pathogens and have been found in spoiled oysters (Cruz-Romero et al., 2008; Yu et al., 2021; Chen et al., 2013). To provide further insights into the metabolic traits of *Pseudomonas* species from the eastern oysters, we conducted whole genome sequencing (WGS), comparative genomic analysis and BIOLOG-based physiological assessments on isolates obtained from the eastern oysters (Pathak et al., 2021). Interestingly, strains that were isolated from the mantle fluid [(*P*. *stutzeri* (MF28)] and tissues [(*P*. *alcaligenes* (OT69)], were metabolically versatile relative to those isolates obtained from the overlaying oyster bed water column [(*P*. *aeruginosa* (WC55)] (Pathak et al., 2022). Specifically, *P*. *aeruginosa* (WC55) preferred polymers over other carbon sources, while *P*. *stutzeri* (MF28) responded better to carbohydrates, while *P*. *alcaligenes* (OT69) showed similar metabolic responses to both polymers and carbohydrates, respectively. Additional studies are required to ascertain whether the eastern oyster associated *Pseudomonas* spp. provide a beneficial service to the host, such as bioremediation of marine pollutants and providing growth promoting vitamins or mediate spoilage via their pathogenic functions.

Though not the main purpose of this study, we also analyzed the archaeal members in the eastern oyster shotgun metagenomes. Archaea are a small fraction of the microbial communities in oysters (Unzueta-Martínez et al., 2022), if not present at all (Pierce and Ward, 2019). Similarly in our study, archaea were represented at only 0.76% in the taxonomic annotation; with dominance of Euryarchaeota (41%) and Thaumarchaeota (24%), respectively (data not shown). The oyster-associated archaeal genera mainly belong to *Methanosarcina* (12%) and *Nitrosopumilus* (10%). Based on the underrepresentation of archaeal communities, it is likely that conditions for the proliferation of this group, such as anaerobiosis, were likely not present in the oysters analyzed in this study.

### 3.2. Alpha and Beta-Diversity Analysis

The bacterial genus level data were statistically evaluated using Shannon’s index for differences potentially driven by seasons and ploidy using alpha and beta diversity analysis. Samples were assigned to 7 different groups based on the collection dates as stated earlier, and some low sequence samples were removed from the analysis. Regardless, this represents a good snapshot of the oyster-associated microbiomes over an annual cycle. Shannon’s diversity, shown in Fig. 2A, was significantly higher at later sampling timepoints but bacterial diversity in diploids and triploids were not different, possibly due to low replication (although p-value for genotypes was 0.056). Alpha diversity analysis revealed that warmer months had increased diversity relative to colder months (Fig 2B). It is widely accepted that warmer temperatures increase marine bacterial abundances (Clare et al., 2022), which goes well with our observation of enhanced bacterial biomass in the oyster samples collected in the warmer season.

**Fig 2.**
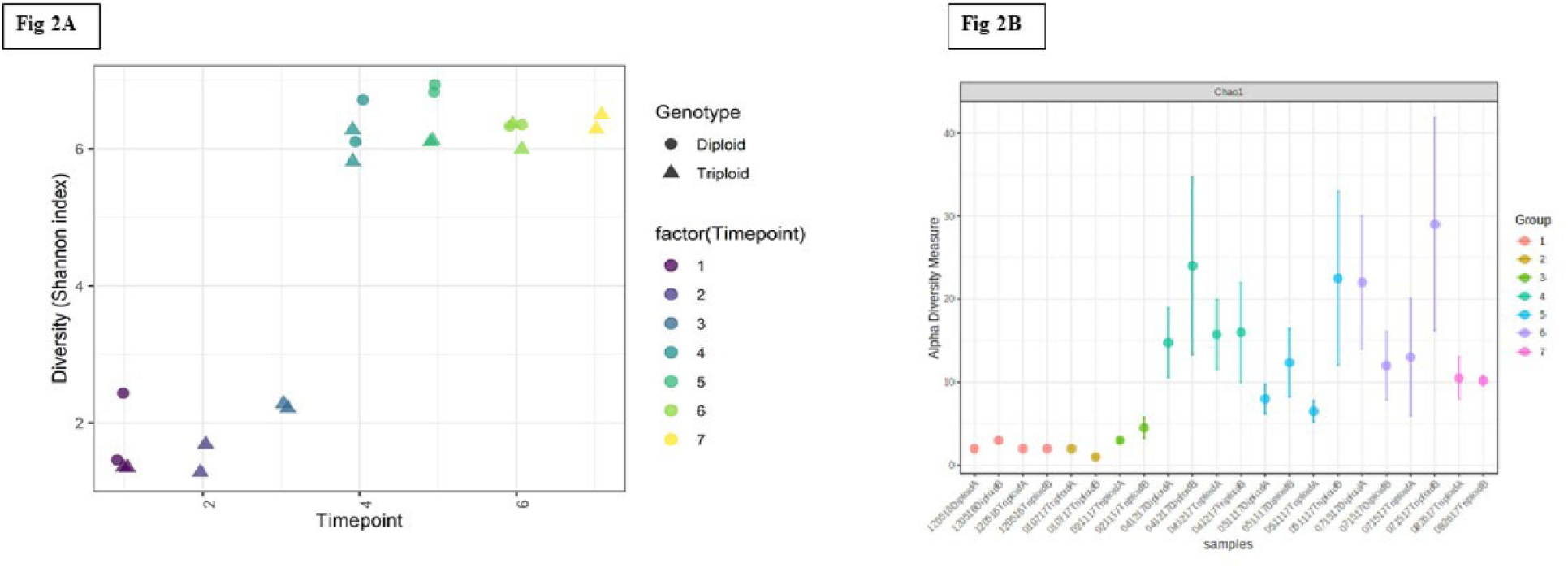
Diversity analysis for each month is represented by a unique color identified in diploid eastern oyster vs. triploid populations, sampled in duplicates (each time point is labeled as A or B), over an annual cycle. Shown are A, Shannon’s diversity analysis, which was significantly higher at later sampling timepoints. No differences between genotypes (diploids and triploids) were found, probably due to low replication, though p-value for genotypes was 0.056; B, alpha diversity calculated from taxonomic data; warmer months were higher in diversity relative to the colder months.

Beta diversity, which is the diversity between different samples and significance testing was also conducted as shown in Fig. 3. Similar to the trend shown in the alpha diversity analysis, beta diversity also revealed that the microbiomes were more similar in oysters sampled in the colder months of December-February than those sampled in warmer months (Fig. 3A and B), as has also been recently shown in Australia (Akter et al., 2023). Microbiomes did not differ between diploid and triploid oysters despite the relatively larger size of the triploid oysters-we noted that diploid oysters ranged from 21-35 mm in height and 1-3 g in weight; triploid oysters heights were 12-27 mm and weights were 0.5-1 g, at the start of the growing period. After a year of growth, diploids were 85-95 mm in height and 54-91 g in weight, and triploids were 71-105 mm in height and 75-110 g in weight. Only few studies are available to compare how ploidy impacts oysters, mostly on mortality, yields, growth rates and susceptibility to diseases (De Decker et al. 2011; Benhaïm et al. 2020). Studies on other seafood, especially fish (Cantas et al., 2011) found that triploids contain significantly higher gut microbiota levels, with increases in *Pseudomonas* spp., *Pectobacterium carotovorum*, *Psychrobacter* spp., *Bacillus* spp., and *Vibrio* spp.

**Fig. 3.**
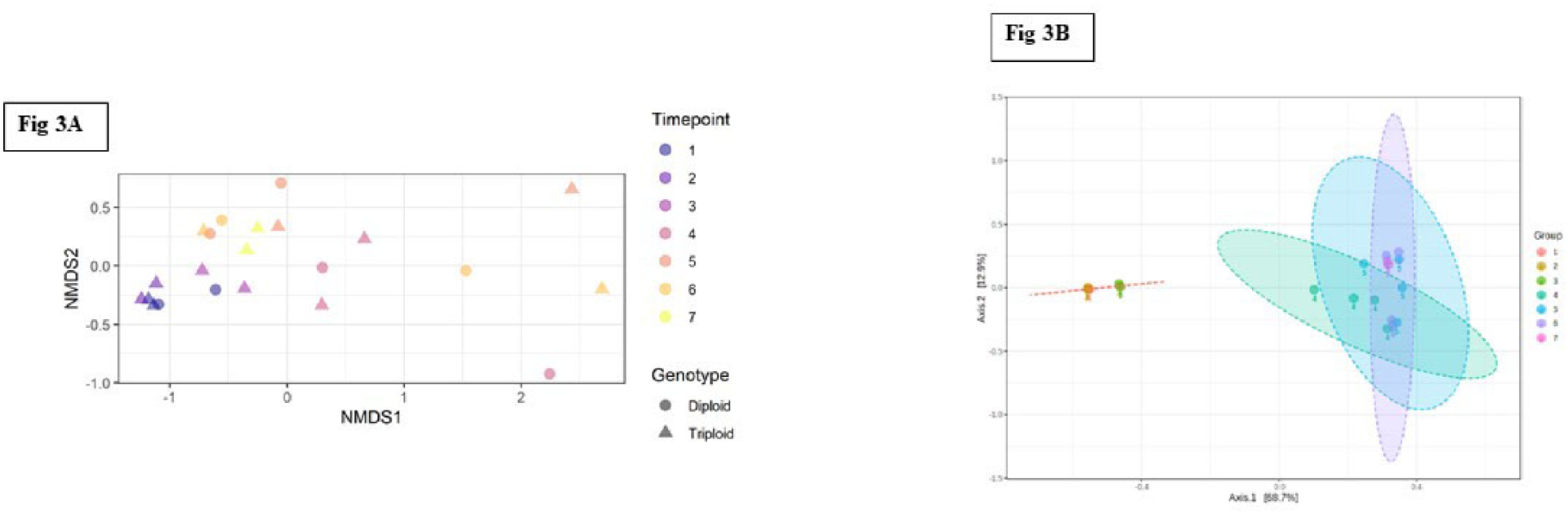
Ordination of bacterial diversity at the genus level, across ploidy and time points which revealed that the oyster microbiomes in colder months of December-February were significantly different to those from warmer months. Shown are A, non-metric multidimensional scaling (NMDS) plot of the beta diversity using Bray-Curtis dissimilarities; B, PCoA PERMANOVA plot of the beta diversity using Bray-Curtis dissimilarities.

### 3.3. Gene Functional Analysis of the Eastern-Oyster Microbiomes Over an Annual Cycle

Functional genes annotated from the oyster shotgun metagenome sequences were organized into the SEED hierarchy (Fig. 4). At broad levels (levels 1 and 2), functional profiles were very similar; smaller differences were found at lower levels (data not shown). Some differences were apparent at subsystems related to bacterial respiration and they tended to increase in later timepoints; a trend similar to the taxonomic assessment, where the latter warmer timepoints revealed a higher abundance and diversity of oyster associated microbiota. The major metabolic functions identified across oyster samples belonged to respiration and metabolism of carbohydrates, RNA, and protein.

**Fig. 4.**
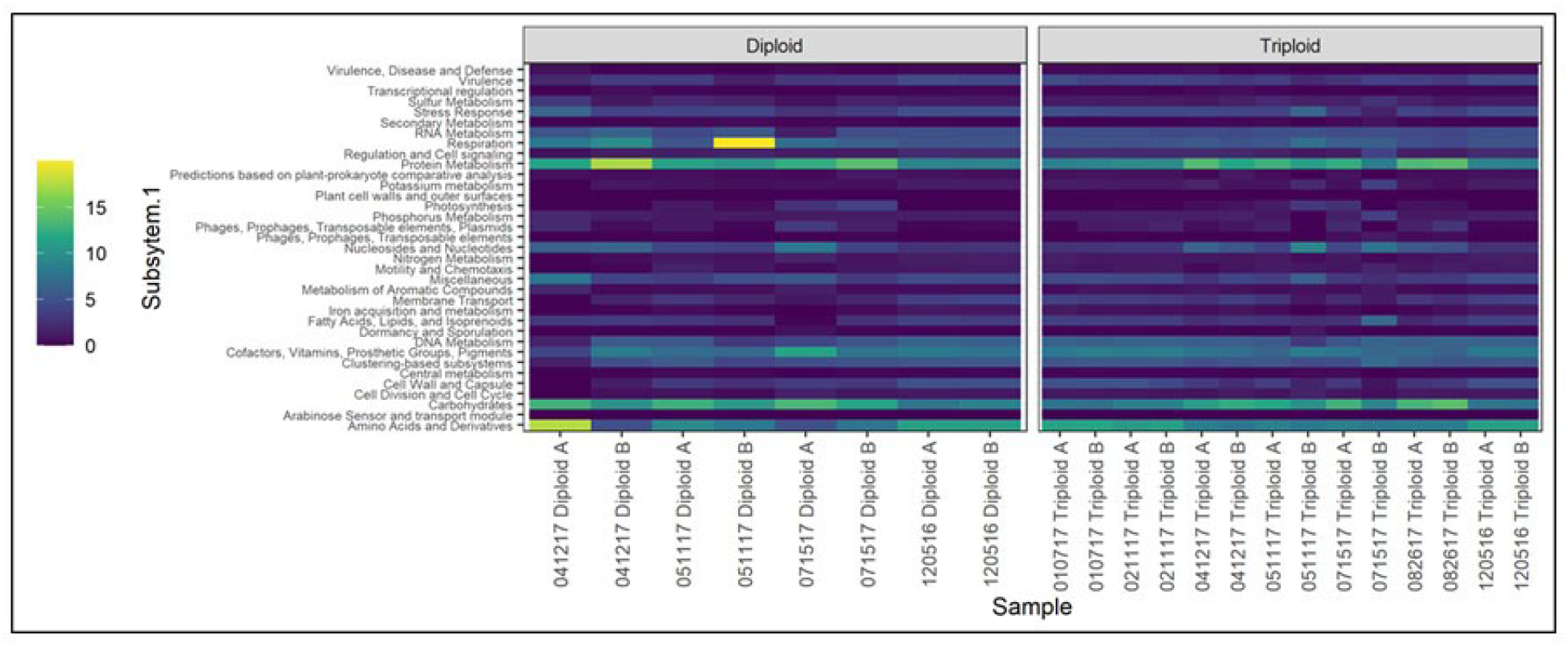
Functional genes annotated from the oyster shotgun metagenome sequences were organized into the SEED functional hierarchy. At broad levels (levels 1 and 2), functional profiles were very similar. Smaller differences were found at lower levels. Some difference were apparent at subsystems related to bacterial respiration, they tended to increase in later timepoints.

We also conducted a permutational multivariate analysis to test for differences in taxonomic composition and gene function over an annual cycle in diploid and triploid oysters (Table SI-1). This showed that taxonomic profiles changed significantly over time, with little or no significant differences observed between genotypes (ploidy). However, post-hoc tests did not find significant differences between timepoints, which can however be due to the low number of replicates. Because we did not find differences between genotypes, we pooled diploids and triploids for the general model; differences among sampling dates explained around 56% of the variation of taxonomic profiles; thus, ploidy was an insignificant factor relative to time-points. Similar to the taxonomic data, no significant differences were found for the gene functional data over the annual time-period in the sampled oysters. Note that these findings are not in line with Sakowski et al., 2020, which showed significant enrichment (p < 0.05) in KEGG based gene functions within the autochthonous “core” microbiomes of the eastern oysters relative to the allochthonous microbiota. Sakowski et al., 2020 also found genes associated with N-cycling, such as dissimilatory nitrate reduction, nitrogen fixation, and nitrification pathways relative to the allochthonous bacterial community.

### 3.4. Differential Abundance of Taxa and Gene Functions

Differential abundance testing of the bacterial taxa (top 20) and genes functions identified at the subsystem level 3 were investigated as a function of two factors-sampling timepoint and genotype (ploidy). As shown in Fig. 5A and B, a general linear model showed several bacterial groups were significantly different when sampling began during the colder months, compared to the latter warmer months, including *Psychrobacter* and *Synechococcus* species-the two most abundant genera identified in this study (Fig. 1A and B). Such seasonal trends in the Pacific oysters and mussels gut microbiomes have recently been published by Akter et al., 2023. Interestingly, *Psychrobacter* was differentially abundant in both diploids as well as triploid oysters, when analyzed separately (Figs. 5B and C). Differential analysis of gene function data revealed several metabolic functions declining at latter timepoints (Fig. 5D), which is counterintuitive because the bacterial taxonomic abundances increased in the mid and latter timepoints, as suggested by diversity analysis (Fig. 2B). Only four gene functions were differentially abundant in diploid oysters (Fig. 5E); conversely, this was not the case in triploids, where several gene abundances were differentially abundant (Fig. 5F). It is unclear as to why more taxa and gene functions were differentially abundant in the triploids relative to the diploid oysters.

**Fig. 5.**
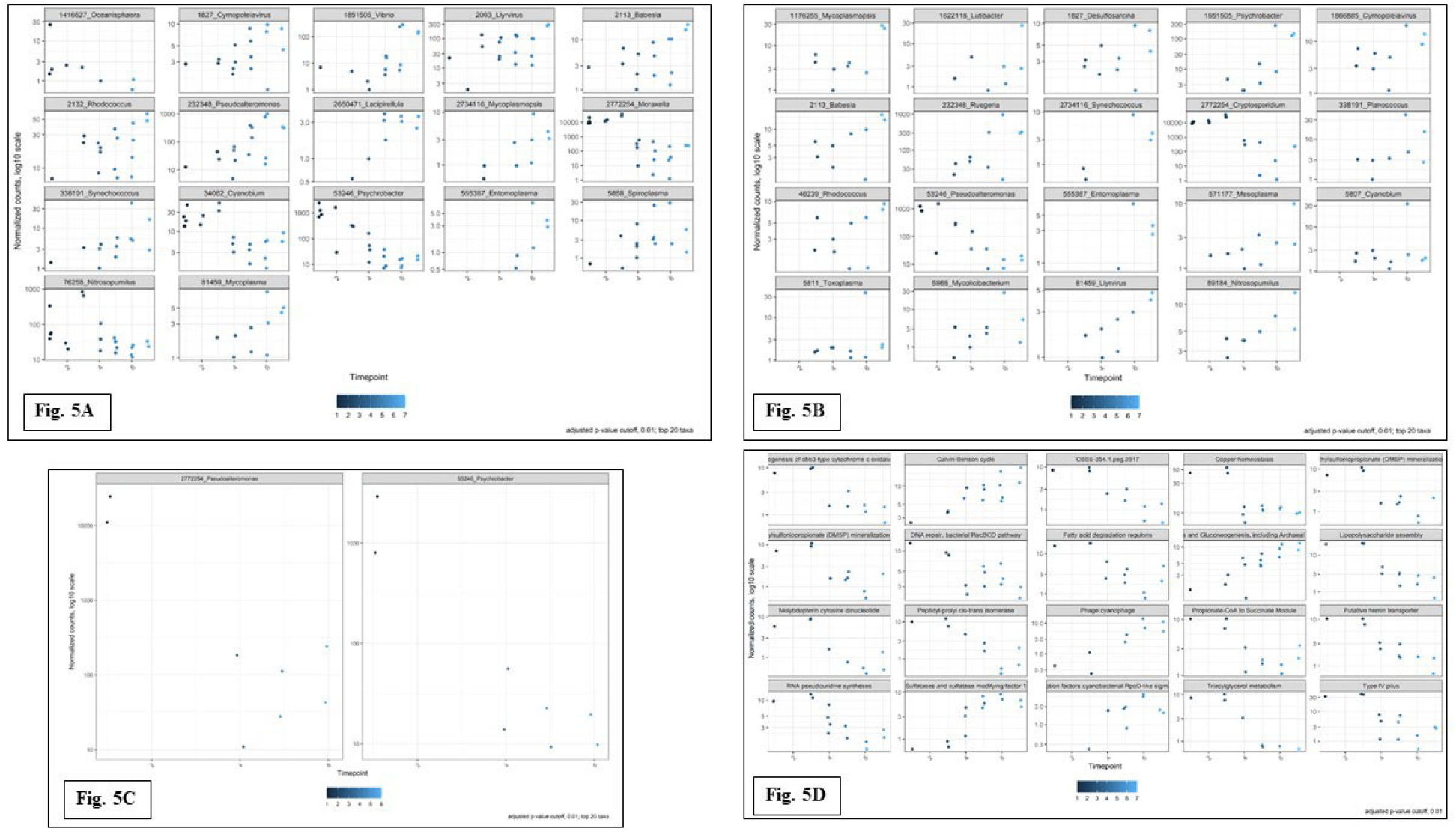
Differentially abundant taxa (genus level) and gene functions (subsystem level 3) identified from the eastern oyster metagenomes over an annual cycle. Shown are A, differentially abundant taxa using a general linear model; B, differentially abundant taxa in diploid oysters; C, differentially abundant taxa in triploid oysters; D, differentially abundant gene functions using a general linear model; E, differentially abundant gene functions in diploid oysters; C, differentially abundant gene functions in triploid oysters, respectively.

### 3.5. Statistical Correlations Between Bacterial Taxa and Gene Functions

The top 10 bacterial taxa at the genus level and gene functions from SEED subsystem level 1 were analyzed statistically (Fig. 6). Dendrogram analysis confirmed that eastern oyster microbiomes clustered more tightly as a function of sampling date and not from differences in ploidy (Fig. 6A). Seven clusters were observed in the dendrogram plot with one outlier (sample from Dec 05 2016 diploids) and ploidy seems insignificant relative to sampling date for bacterial community structure in this study. The Non-metric Multi-Dimensional Scaling plots (NMDS) (Fig. 6B) at a similarity at 75% and 95% revealed interesting correlations between bacterial top 10 genera and SEED subsystem level 1 features. Specifically, at the 95% similarity threshold, *Psychrobacter,* a member of the “core” microbiome regardless of date or ploidy status, correlated strongly to gene functions of central metabolism, DNA metabolism and carbohydrates (Fig. 6B). Other taxa that were within the 95% similarity index threshold were *Moraxella*, *Pseudoalteromonas*, *Brochothrix*, *Halomonas*, and these correlated to gene functions of cofactors, vitamins, prosthetic groups, pigments; cell division and cell cycle; cell wall and capsule; amino acids and derivatives; dormancy and sporulation, and clustering-based subsystems. The other taxa that correlated at the 95% threshold were *Vibrio* and Acinetobacter with no affiliation to any of the top 10 gene functions. *Pseudomonas*, *Shewanella* and *Flavobacterium* were not significantly correlated to other taxa or gene functions (Fig. 6B). At a threshold of 75% similarity, all the above stated top 10 bacterial genera and gene functions were correlated. Based on this, it is likely that *Psychrobacter* species perform significant functional roles in the eastern oyster microbiome and thus, further studies to assess the role of this genus within the broader context of the oyster microbiome and how it intersects with oyster health and productivity are thus warranted.

**Fig. 6.**
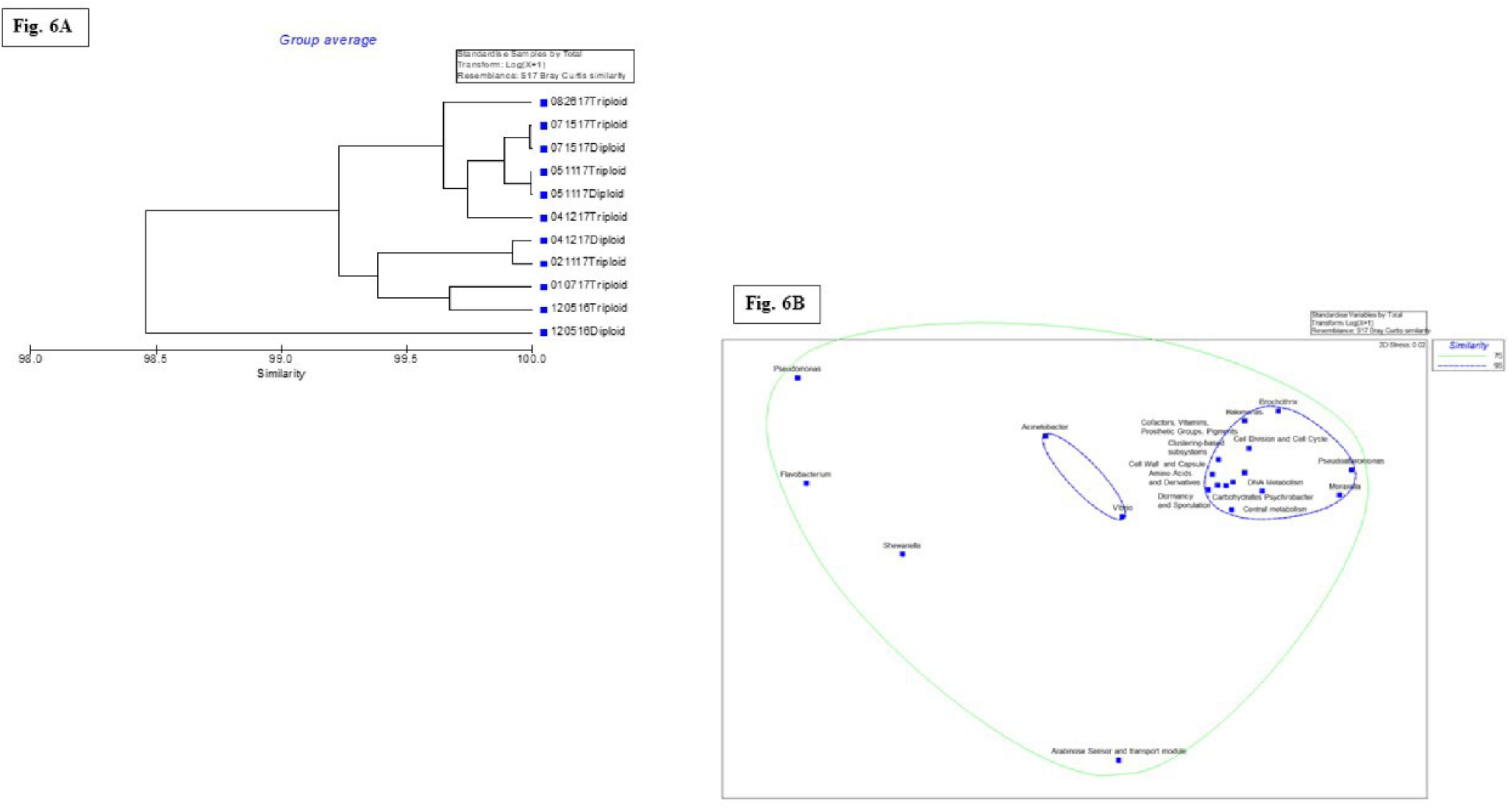
Statistical correlations between the top 10 bacteria (genus level) and SEED subsystem (level 1) is shown plotted as a dendrogram analysis (A) or a Non-metric Multi-Dimensional Scaling plots (NMDS) (B), respectively, where similarity at 75% and 95% is shown.

### 3.6. Metagenome Assembly, Phylogenetic, and ANI Analysis

To better characterize the dominant species, metagenome assembly was performed, followed by phylogenetic analysis of select MAGs (metagenome-assembled genomes). Eleven MAGs were obtained, six of which remained after filtering based on completeness as measured using BUSCO (Table 1A). Four of the six were classified as *Psychrobacter* (Fig. 7), one as *Pseudoalteromonas* and one as belonging to the same family as COTS27, a bacterial symbiont of the crown-of-thorns starfish (Wada et al., 2020). The MAGs consist of many contigs (M = 591.5, SD = 213.4) and exhibit a modest level of completeness (M = 51.6% SD = 22.2%). The MAG SPAdes-group-0.1.fa displays a higher level of duplicated genes (8.1%), which could reflect the presence of genes from more than one species or issues with the assembly. Phylogenetic analysis of inferred protein sequences was performed to compare the *Psychrobacter* MAGs to each other and to *Psychrobacter* RefSeq assemblies available in the NCBI Assembly database. The resulting maximum likelihood tree supports that the four *Psychrobacter* MAGs are each most closely related to separate *Psychrobacter* RefSeq assemblies, and not part of a single lineage.

**Table 1A.** Metagenome-assembled genomes (MAG) based taxonomy and completeness from the eastern-oyster derived shallow shotgun metagenomic data.

| MAG | Taxonomy (family;<br>genus) | Contig<br>number | Total length bp | BUSCO % complete (single<br>copy, duplicated,<br>fragmented) |
| --- | --- | --- | --- | --- |
| SPAdes-group-0.1.fa | *Moraxellaceae*;<br>*Psychrobacter* | 893 | 3004744 | 41.2 (22.6, 8.1, 10.5) |
| SPAdes-group-0.4.fa | *Moraxellaceae*;<br>*Psychrobacter* | 474 | 1479441 | 38.7 (30.6, 0, 8.1) |
| SPAdes-group-0.5.fa | *Moraxellaceae*;<br>*Psychrobacter* | 455 | 1873903 | 47.6 (38.7, 0, 8.9) |
| SPAdes-group-0.7.fa | *Alteromonadaceae*;<br>*Pseudoalteromonas* | 745 | 4536590 | 77.4 (75.8, 0.8, 0.8) |
| SPAdes-group-0.8.fa | *Moraxellaceae*;<br>*Psychrobacter* | 665 | 1695251 | 25 (16.1, 0, 8.9) |
| SPAdes-group-0.11.fa | COTS27; _ | 317 | 1427112 | 79.9 (70.2, 0, 9.7) |

**Fig. 7.**
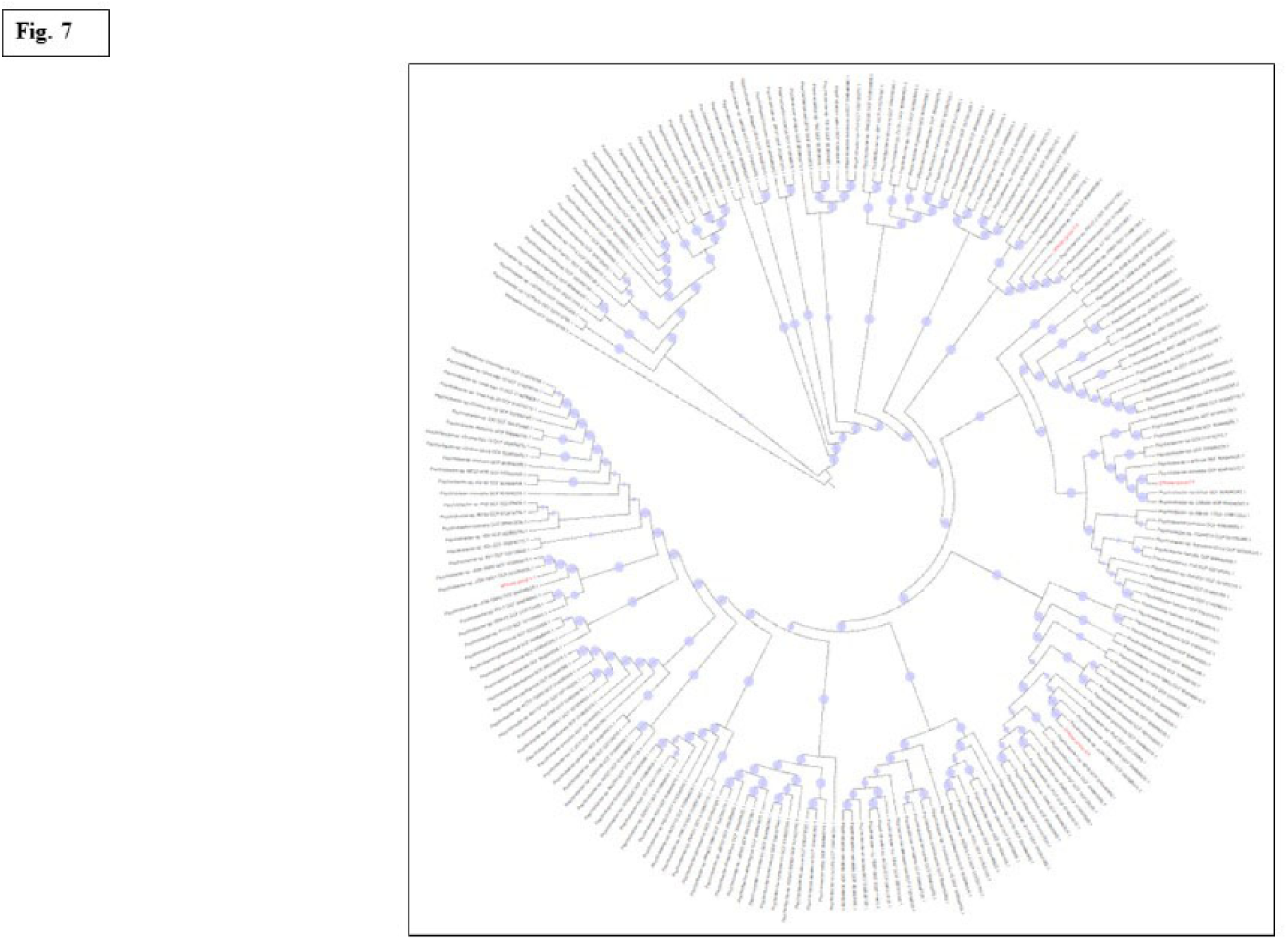
Maximum likelihood tree of *Psychrobacter* MAG proteomes obtained from this study is shown (red labels) relative to RefSeq *Psychrobacter* proteomes from NCBI (black labels). Bootstrap support from 100 replicates is indicated using blue circles.

To complement the phylogenetic analysis, whole-genome average nucleotide identity (ANI) was calculated between each *Psychrobacter* MAGs and all *Psychrobacter* RefSeq genomes (Table 1B). MAGs SPAdes-group-0.4.fa and SPAdes-group-0.5.fa exhibit high ANI (98%) with their respective matches, and the similarity involves most of the MAG sequence (97%). These ANI values are characteristic of intra-species relationships (Jain et al., 2018). SPAdes-group-0.8.fa also showed high ANI with its top match (97%) but the similarity involved just 71% of the MAG. SPAdes-group-0.1.fa exhibited a lower ANI (88%) with its top match, which is lower than the intra-species level (>95%) (Jain et al., 2018), and the similarity involved 71% of the MAG. Together these results further support the presence of multiple *Psychrobacter* species within the eastern-oyster microbiome. Additional metagenomic sequencing to provide more complete, higher-quality assemblies should help further resolve the relationships between these potentially novel isolates or species and existing *Psychrobacter* isolates and help clarify their potential role(s) in the oyster holobiont.

**Table 1B.** Top ANI matches for *Psychrobacter* MAGs obtained from the eastern-oyster derived shallow shotgun metagenomic data.

| MAG | Highest ANI | Percentage of MAG aligned as orthologous matches | RefSeq organism name | RefSeq assembly accession |
| --- | --- | --- | --- | --- |
| SPAdes-group-0.1.fa | 88.03 | 70.82 | *Psychrobacter* sp. KH172YL61 | GCF_007182895.1 |
| SPAdes-group-0.4.fa | 98.32 | 97.22 | *Psychrobacter* sp. JCM18903 | GCF_904846635.1 |
| SPAdes-group-0.5.fa | 98.45 | 97.08 | *Psychrobacter celer* | GCF_904845945.1 |
| SPAdes-group-0.8.fa | 96.61 | 70.58 | *Psychrobacter maritimus* | GCF_904846345.1 |

### 3.7. Denitrification Genes in *Psychrobacter* MAGs and Similar RefSeq Genomes

The denitrification process is performed by a diverse range of microorganisms, especially under oxygen depleted conditions, with the complete reduction of NO^3-^ to N^2^ performed via a four-step process, involving the membrane-bound NO^3-^reductases (narG) and/or the periplasmic napA genes, the NO^2-^ reductases NirK (copper-containing) or NirS (containing cytochrome cd1) encoded by nirK and nirS; the NO reductases cNor (cytochrome c dependent) or qNor (quinol-dependent) encoded by cnor and qnor; and the N_2_O reductase NosZ.

Sequence-based identification of denitrification genes in the *Pyschrobacter* MAGs indicates the presence of *norB* in SPAdes-group-0.8.fa (Table 1C). Given the incomplete nature of the MAGs, the search for denitrification genes was extended to the top ANI match for each MAG, under the assumption that the more complete RefSeq genomes would provide a more comprehensive view of the denitrification gene repertoire. This search of the highly similar genomes revealed the presence of *narH* and *narI* for RefSeq genome GCF_904846635.1 (the top ANI match for SPAdes-group-0.4.fa), and the presence of *narH*, *narI*, *nirK*, and *norB* for RefSeq genome GCF_904846635.1 (the top ANI match for SPAdes-group-0.8.fa). Other denitrification genes, especially for complete denitrification, such as nosZ could not be identified based on the MAG approach. Deeper sequencing depths and coverages will likely enhance our understanding on the metabolic repertoire of *Psychrobacter* spp., associated with eastern oysters.

**Table 1C.**
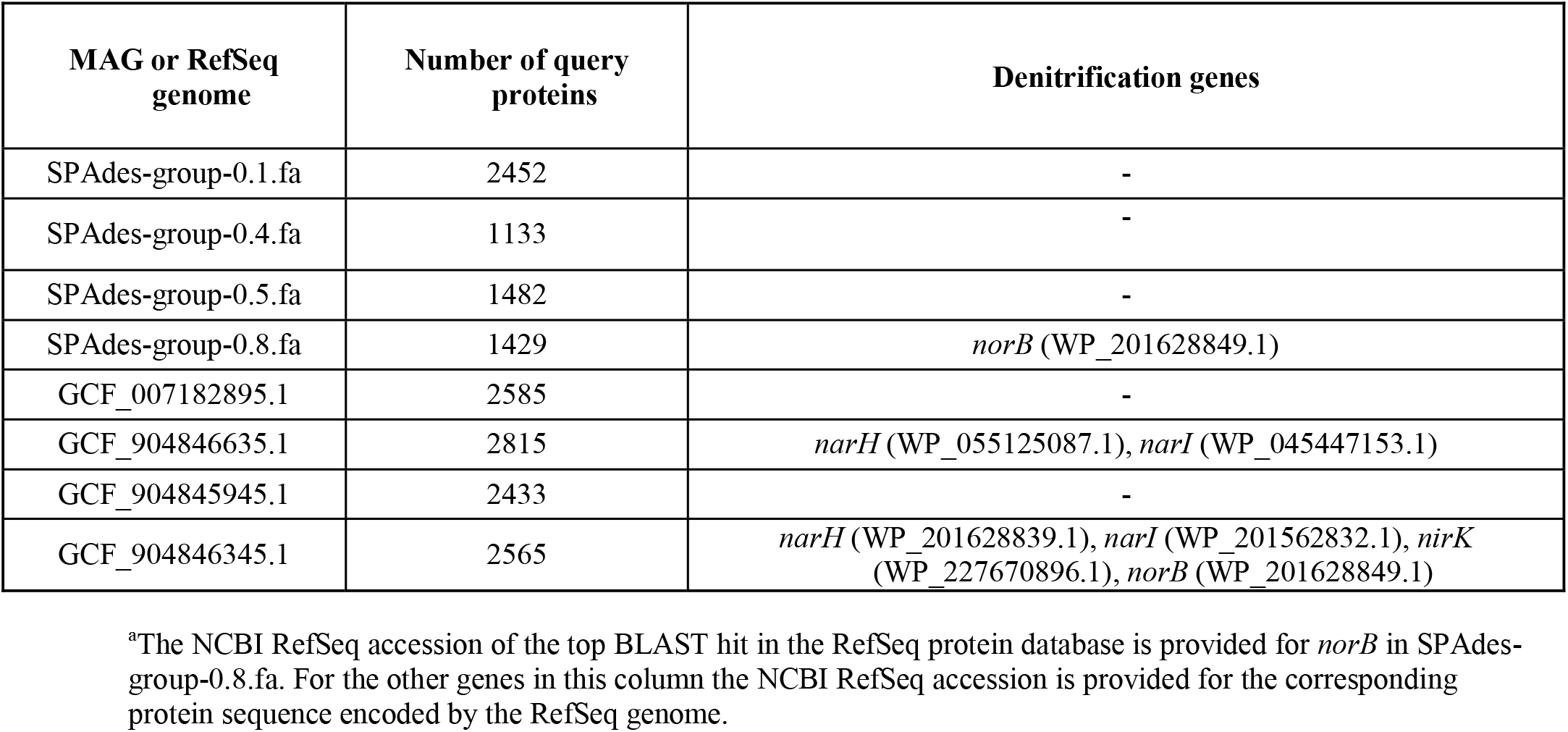
Denitrification genes identified in *Psychrobacter* MAGs obtained from the eastern-oyster derived shallow shotgun metagenomic data and similar RefSeq genomes.

### 3.8. Quantitative Microbial Elemental Cycling (QMEC) Analysis

When QMEC was used on genomic DNA extracted from homogenized diploid and triploid oysters, a total of 71 genes responsible for CNPS cycling amplified successfully. Note that we did not separate the bacterial biomass from the oyster tissue homogenates, which likely caused PCR inhibition from the overrepresented oyster DNA/ mucous and consequently, gene copy abundance/g tissue from only 8 samples covering 3 sampling timepoints yielded usable data (Figs. 8A-D). Regardless, this is the first application of QMEC to understanding biogeochemical cycling in eastern oysters. Relative abundances of CNPS genes were higher in samples collected in December 2016 but did not vary much except in samples collected on April 12, 2017, which seems to be an outlier. PCA conducted on absolute and relative CNPS gene copy numbers revealed differences between diploid and triploid samples but not the sampling time points (Fig. 8E-F), which was not in line with the taxonomic data-which suggested that season but not ploidy was the stronger of the two factors in structuring oyster microbiomes. However, it may not be prudent to compare shotgun and qPCR data.

**Fig. 8.**
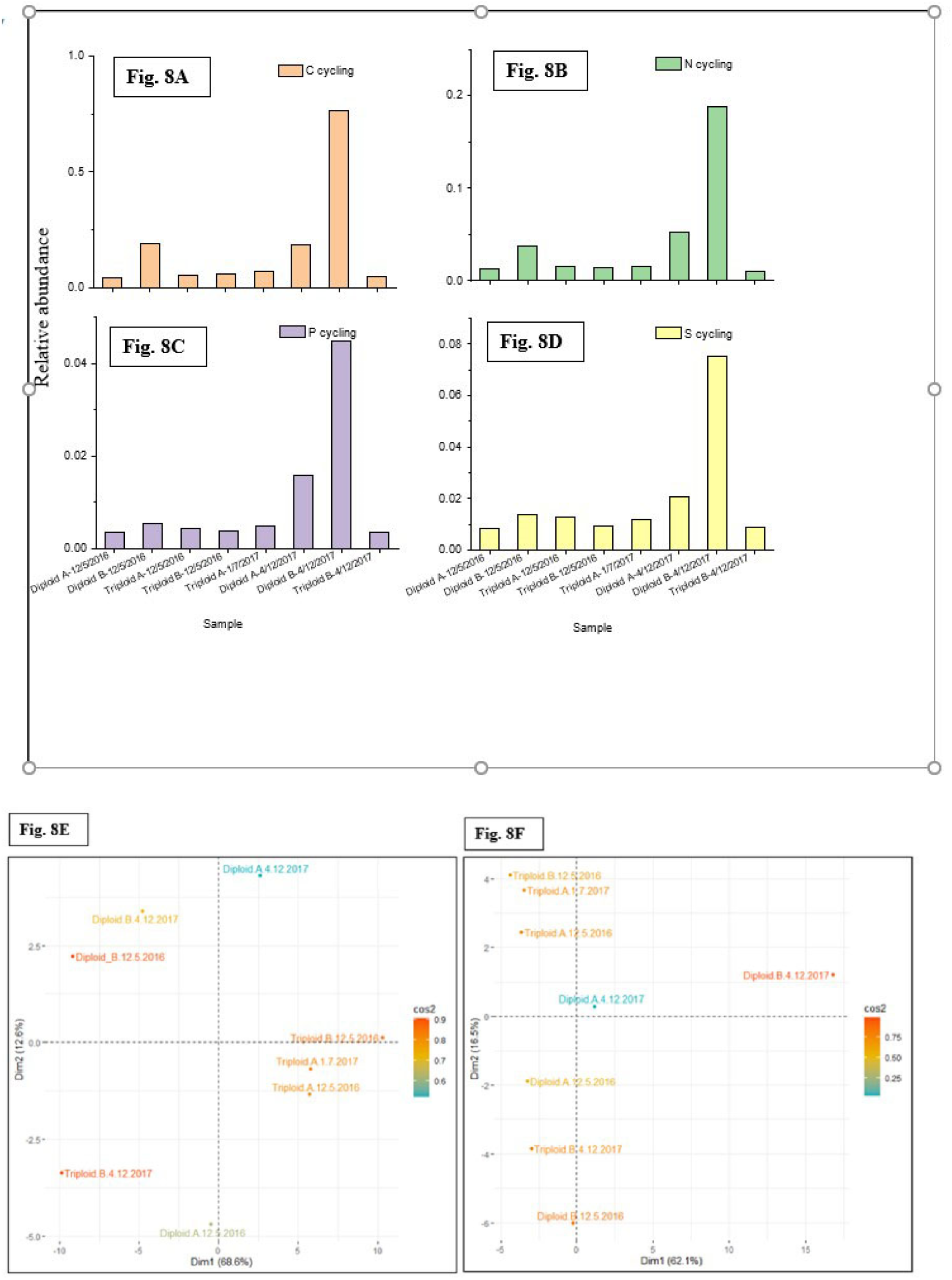
Shown are A, relative abundance of carbon (C); B, nitrogen (N), C, phosphorus (P) and sulfur (S) cycling genes using QMEC analysis. Also shown are principal component analysis (PCA) of absolute (E) and relative (F) gene copy numbers that indicated ploidy to have a significant impact relative to sampling time points.

Among the carbon cycling genes, C-fixation was the most abundant process followed by methane production and oxidation (Fig. 9A). Carbon fixation has been found to be an enriched gene function in the mucus microbiomes associated with corals (Badhai et al., 2016) as well as the microbiome of the Black-Lipped Pearl Oyster (Dubé et al., 2019). *Psychrobacter* spp. were abundant in the coral mucus study and it is likely that they are important in carbon cycling processes (McKew et al., 2012; Badhai et al., 2016). Methane formation (methanogenesis) and oxidation has also been shown to be driven by macrofaunal organisms, such as the bivalve *Limecola balthica* and the polychaete *Marenzelleria arctia* (Bonaglia et al., 2017). Furthermore, symbiotic associations between methane-producing archaeal communities and bivalves have also been previously reported (Bonaglia et al., 2017); these processes were also identified in the tested eastern oysters, albeit at lower levels. Among the tested 22 nitrogen cycling genes, denitrification was the highest followed by ammonification (Fig. 9B). Note that denitrification in oysters can contribute significantly in N transformations in pelagic regions (Jiang et al. 2020), as well as in oyster reefs, aquaculture sites, and their sediments (Hoellein et al. 2015; Mortazavi et al. 2015; Smyth et al. 2016). In fact, it is the oyster-associated denitrifier microbiota that contribute to cycling reactive N into N_2_ gas (Ayvazian et al. 2021; Arfken et al., 2017) via three pathways: (1) through increasing organic matter deposits in sediments, (2) hosting denitrifying bacteria within gills and/ or digestive organs and/or (3) facilitating habitat for other filter-feeding macrofaunal communities (Ayvazian et al. 2021). Organic phosphorus (P) mineralization and sulfur (S) oxidation were the major pathways in P and S biogeochemical cycling processes (Fig. 9C and D).

**Fig. 9.**
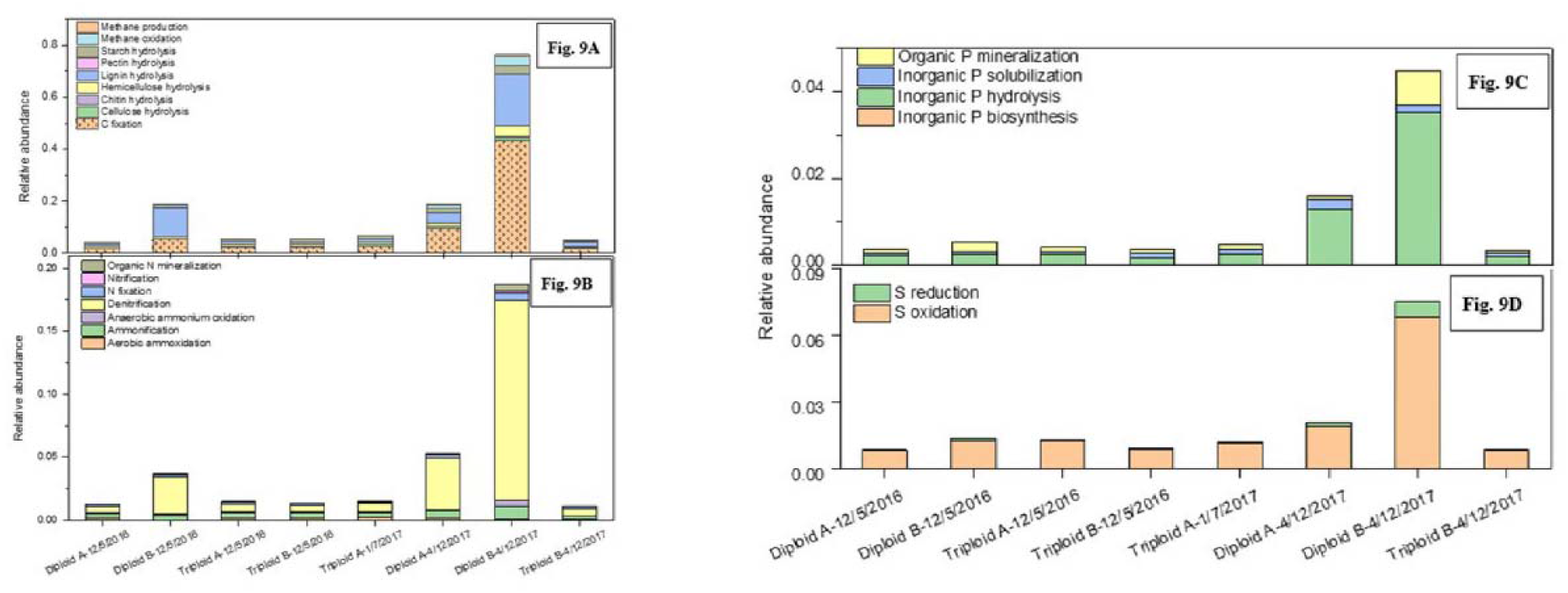
Shown are A, carbon (C); B, nitrogen (N), C, phosphorus (P) and D, sulfur (S) cycling relative gene abundances when QMEC analysis was performed on the diploid and triploid eastern oysters.

Among the denitrification genes encoding for nitrite reductase (nirS or nirK) or nitrous oxide reductase (nosZ), *nosZ*II was the most dominant (Fig. 10A). Under anaerobic conditions, denitrification occurs as a stepwise reduction of nitrate (NO3-) to either nitrous oxide (N_2_O) or dinitrogen gas (N_2_). Complete denitrification takes place via the reduction of nitric oxide (NO) to N_2_O and then the reduction of N_2_O to N_2_ gas. The two microbially-mediated enzymes facilitating the denitrification process are nitric oxide reductase (NOR) and nitrous oxide reductase (N2OR). Based on genotypic studies, nosZ gene has been subdivided into clade I (nosZI) or clade II (nosZII) (Sanford et al., 2010; Jones et al., 2013), with the latter dominant in most habitats including eastern oysters. Clade I nosZ gene containing bacteria mainly belong to Alphaproteobacteria, Betaproteobacteria and Gammaproteobacterial, but clade II nosZ gene has been identified from a variety of bacterial taxa, such as Deltaproteobacteria, Bacteroidetes, and Gemmatimonadetes (Sanford et al. 2012; Jones et al. 2013; Hallin et al. 2018), respectively.

**Fig. 10.**
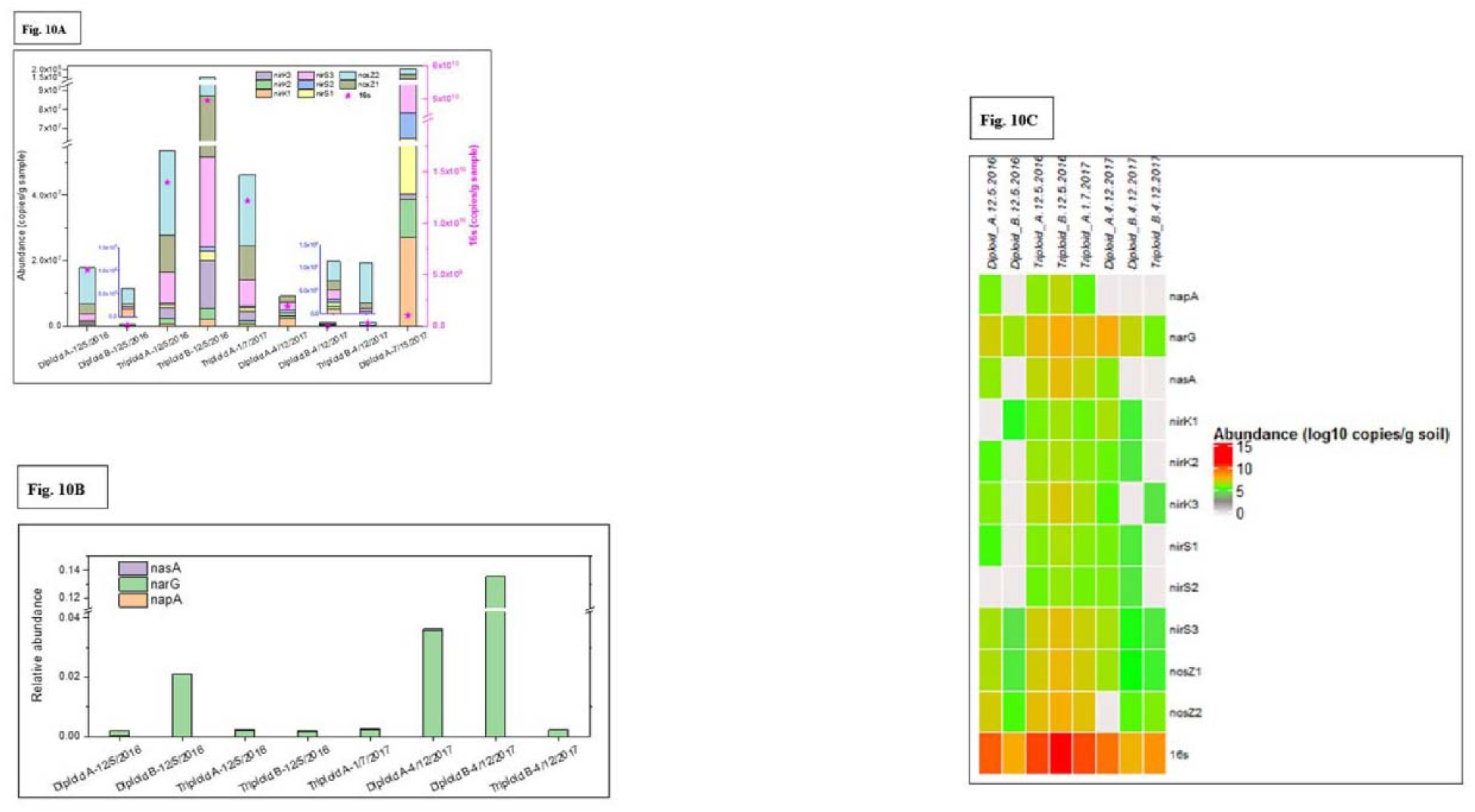
Shown are copy number of specific nitrite and nitrate reductase denitrification genes such as A, nirK, nirS, nosZ; B, napA, narG, nasA, when QMEC analysis was performed on the diploid and triploid eastern oysters and C, heatmap of denitrification genes plotted along with th 16S for comparisons.

More recently, it was found that autochthonous “core” microbiomes within eastern oysters were significantly (p < 0.05) enriched with 933 KEGG based gene functions relative to the 700 gene functions associated with the allochthonous oligotypes (Sakowski et al., 2021). The autochthonous oligotypes were also significantly enriched for several N-cycling genes, such as for dissimilatory nitrate reduction, nitrogen fixation, and nitrification pathways relative to the allochthonous oligotypes. To this end, Arfken et al., 2017 showed that an increase in denitrification rates correlated positively to an increase in the gene abundances of nosZI, but this trend did not hold for the more dominant nosZII. This suggests that oyster-associated microbes possessing the nosZI genes may be more significant to denitrification processes than those that possess the nosZII genes. However, other reports have shown that nosZII denitrifiers are dominant over the nosZI denitrifiers in a variety of different ecosystems (Jones et al., 2013; Orellana et al., 2014), including the eastern oysters (Arfken et al., 2017). This could be because the nosZII gene renders ecophysiological diversity relative to the nosZI gene (Sanford et al., 2012), suggesting that denitrifier microbiotain oysters are likely far more diverse than previously known. Moreover, in this study, the *nasA*, *narG*, and *napA* genes were also observed (Fig 10B) with *narG* dominant. *NarG* encodes for membrane-bound nitrate reductase (Smith et al, 2007), which has previously been found to be high in sediments from oyster farms (Mara et al. 2021) and appears to be dominant in eastern oysters, regardless of ploidy. The higher abundances of nosZII and narG, relative to 16S genes in eastern oysters using the QMEC method are shown in Fig. 10C.

Overall, this study coupled the use of shallow shotgun metagenomics and high-throughput qPCR (HT-qPCR) to enhance our understanding on the resident “core” or autochthonous microbial communities within the eastern oysters. We unequivocally show that the *Psychrobacter* genus is likely one representative of the eastern oyster’s “core” microbiome due to the predominance and stable numbers of this psychotrophic bacteria over an annual growth cycle. Note however that these findings are largely based on shallow shotgun sequencing and future studies conducted using deeper sequencing will likely lead to a comprehensive understanding of the eastern oyster’s “core” microbiomes, thus paving the way for improving oyster productivity and their ecosystem functions via manipulation of their microbial communities. We also plan on conducting additional studies to delineate whether *Psychrobacter* species are beneficial or detrimental to oyster health and associated ecosystem services, including biogeochemical cycling of nutrients. Given the central role microbiomes play in oysters and other bivalve species, it is imperative to establish a universally accepted standard in defining the “core” (Shade and Handelsman, 2012), and by inference, the resident or autochthonous bacterial species for the eastern oysters; studies in this direction are also thus warranted.

## Acknowledgments

This work was mainly supported by a National Science Foundation (NSF) award (#1901371). Other funding also partly supported this research, including Cooperative Agreement NA11SEC4810001 between the National Oceanic and Atmospheric Administration’s Educational Partnership Program for Minority Serving Institutions and the Environmental Cooperative Science Center at Florida A&M University, NSF award #2200615 and three US Department of Energy (DOE) projects, that included the Minority Serving Institution Partnership Program (MSIPP) (task order agreements #0000403081, #0000403082, and #0000456318). We thank B. Ballard and A. Wynn form the Wakulla Environmental Institute for supplying oysters and assisting with field collections. Technical help provided by Dr. Meenakshi Agarwal, Dr. Rajneesh Jaswal, Mr. Rajesh S. Rathore, is appreciated.

## Author’s Contribution

AC, AP, CHJ, MM contributed to conception and design of the study. AP organized the database and performed the statistical analysis. AP wrote the first draft of the manuscript. AC, AP, MM, PS, JQS, YZ, XYZ, CC and CHJ wrote sections of the manuscript. All authors contributed to manuscript revision, read, and approved the submitted version.

---

**Fig. SI-1:**
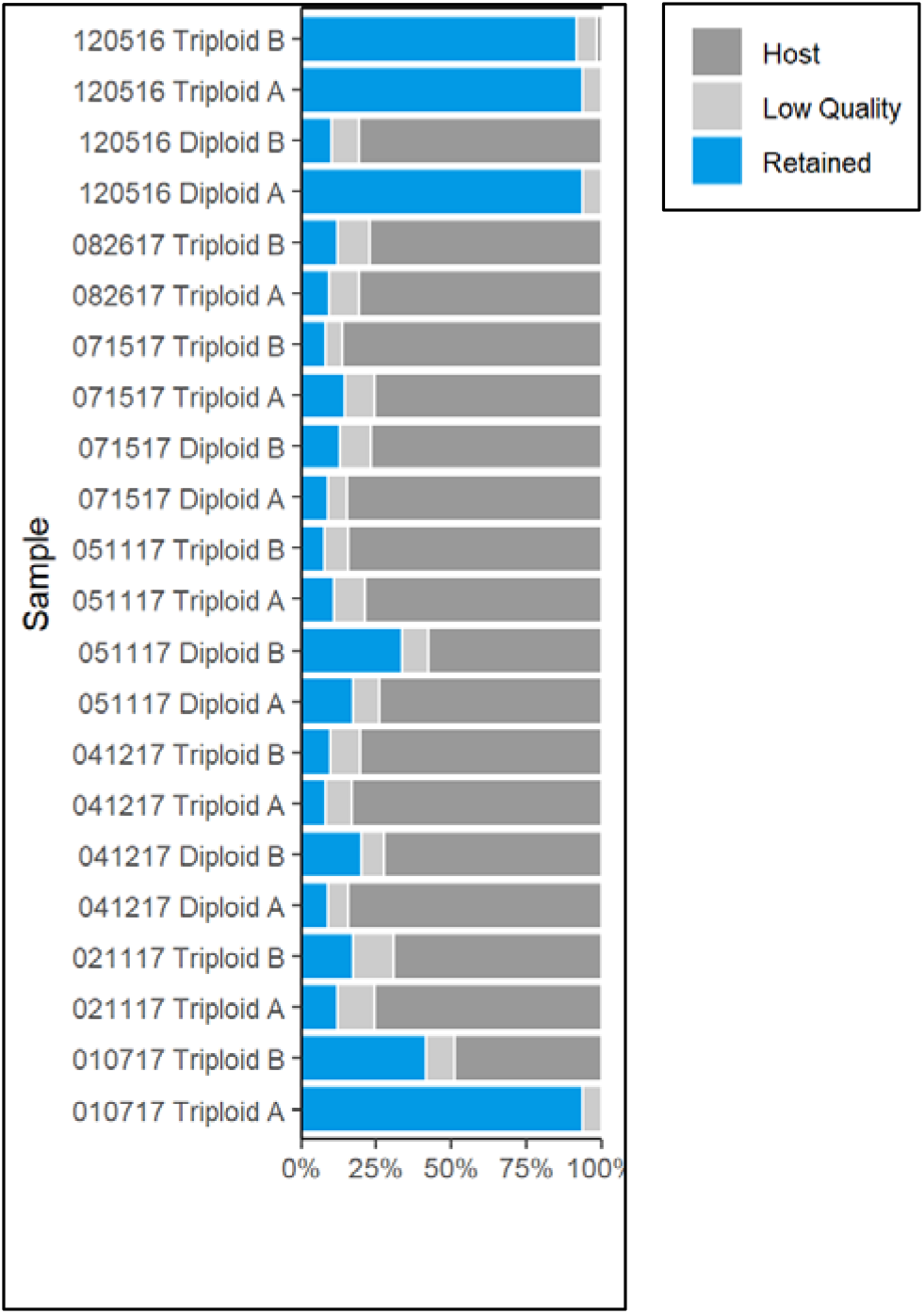
A collection of over 113.2 million raw reads and 98.3 quality-filtered reads were obtained from the eastern oysters collected from oyster bay, FL. Sequencing was performed on duplicate oyster samples at each time point, labeled as A and B and each sample name begins with the date they were collected. Around 34.3 (34.8%) and 0.5 million (0.51%) of the quality-filtered reads were assigned to the oyster host (*Crassostrea virginica* genome v3.0 (Accession number GCF_002022765.2) and the human genome (Genome Reference Consortium Human Reference 37), which were removed prior to further processing.

**Table SI-1:** Analysis of Variation. We used a Permutational multivariate analysis variation to estimate the effects of experimental factors on both taxonomic and functional profiles. We found that taxonomic profiles changed significantly over time but there were no significant differences between genotypes (ploidy). Post-hoc test did not find significant difference between timepoint pairs after multiple test correction, likely due to the low number of replicates. In both diploid and triploid samples, we also found differences mainly due to timepoints. Since we did not find differences between genotypes we used all the samples together for the general model. Timepoint differences explained around 56% of the variation of taxonomic profiles. No significant differences were found for the functional profiles.

**PERMANOVA taxonomic profiles general model**
|  | Df | SumOfSqs | R2 | F | Pr..F. |
| --- | --- | --- | --- | --- | --- |
| Genotype | 1 | 0.226 | 0.045 | 2.071 | 0.140 |
| Timepoint | 1 | 2.837 | 0.560 | 25.948 | 0.001 |
| Genotype:Timepoint | 1 | 0.031 | 0.006 | 0.284 | 0.886 |
| Residual | 18 | 1.968 | 0.389 | NA | NA |
| Total | 21 | 5.062 | 1.000 | NA | NA |

**PERMANOVA taxonomic profiles diploid samples**
|  | Df | SumOfSqs | R2 | F | Pr..F. |
| --- | --- | --- | --- | --- | --- |
| Timepoint | 1 | 0.928 | 0.544 | 7.169 | 0.01 |
| Residual | 6 | 0.776 | 0.456 | NA | NA |
| Total | 7 | 1.704 | 1.000 | NA | NA |

**PERMANOVA taxonomic profiles triploid samples**
|  | Df | SumOfSqs | R2 | F | Pr..F. |
| --- | --- | --- | --- | --- | --- |
| Timepoint | 1 | 1.940 | 0.62 | 19.54 | 0.001 |
| Residual | 12 | 1.191 | 0.38 | NA | NA |
| Total | 13 | 3.131 | 1.00 | NA | NA |

**PERMANOVA functional profiles general model**
|  | Df | SumOfSqs | R2 | F | Pr..F. |
| --- | --- | --- | --- | --- | --- |
| Genotype | 1 | 0.099 | 0.047 | 0.751 | 0.703 |
| Timepoint | 1 | 0.174 | 0.082 | 1.320 | 0.215 |
| Residual | 14 | 1.849 | 0.871 | NA | NA |
| Total | 16 | 2.122 | 1.000 | NA | NA |

PERMANOVA functional profiles diploid samples
|  | Df | SumOfSqs | R2 | F | Pr..F. |
| --- | --- | --- | --- | --- | --- |
| Timepoint | 1 | 0.102 | 0.107 | 0.599 | 0.69 |
| Residual | 5 | 0.852 | 0.893 | NA | NA |
| Total | 6 | 0.954 | 1.000 | NA | NA |

PERMANOVA functional profiles triploid samples
|  | Df | SumOfSqs | R2 | F | Pr..F. |
| --- | --- | --- | --- | --- | --- |
| Timepoint | 1 | 0.112 | 0.105 | 0.938 | 0.447 |
| Residual | 8 | 0.957 | 0.895 | NA | NA |
| Total | 9 | 1.069 | 1.000 | NA | NA |

